# Behavioral effects of ethanol in the Red Swamp Crayfish (*Procambarus clarkii*)

**DOI:** 10.1101/2021.06.05.447220

**Authors:** Arnold Gutierrez, Kevin M. Creehan, Giordano de Guglielmo, Amanda J. Roberts, Michael A. Taffe

**Author notes:** Address Correspondence to: Dr. Michael A. Taffe, Department of Psychiatry, 9500 Gilman Drive;, University of California, San Diego, La Jolla, CA 92093; USA.

## Abstract

Alcohol abuse and dependence remains one of the primary preventable sources of human mortality in the United States. Model laboratory species can be used to evaluate behavioral, neuropharmacological and other biological changes associated with alcohol exposure and to identify novel treatment modalities. This report describes methods for evaluating the behavioral effects of ethyl alcohol (ethanol; EtOH) in a crayfish model. Crayfish (*Procambarus clarkii*) were immersed in ethanol baths with concentrations ranging from 0.1 to 1.0 molar, and for durations of 10-30 minutes. Studies evaluated hemolymph alcohol concentration, locomotor behavior in an open field and anxiety-like behavior using a Light/<u>Dark</u> transfer approach. EtOH immersion produced dose dependent increases in hemolymph EtOH concentration (up to 249 mg/dL) and reductions in open field locomotor behavior that depended on EtOH concentration or exposure duration. Under baseline conditions, crayfish exhibit avoidance of the open parts of the locomotor arena and a preference for the covered portion, when available. Acute EtOH immersion decreased time spent in the covered portion of the Light/Dark arena, consistent with a decrease in anxiety-like behavior. EtOH immersion once per day for 5 days did not alter the acute locomotor behavioral response, however increased activity was observed 3 days after the repeated EtOH regimen. Overall, this study shows that this inexpensive, easily maintained species can be used for behavioral pharmacological experiments designed to assess the acute and repeated effects of EtOH.

## 1. Introduction

About 14.4 million adults have an Alcohol Use Disorder (AUD) and some 88,000 Americans die from alcohol-related causes annually, making it the third leading preventable cause of death (https://www.niaaa.nih.gov/sites/default/files/AlcoholFactsAndStats.pdf). It continues to be a pressing concern to learn more about the biological impact of alcohol exposure, so as to generate novel avenues for remediation. A recent review of animal models of substance use disorder calls in part for expansion of behavioral models suitable for substance abuse investigations (Smith, 2020), including invertebrate models. The crayfish, a Genus of numerous species, is a potential animal model which offers significant advantages in terms of cost over more common small vertebrate species such as rats or mice. Species such as the red swamp crayfish (*Procambarus clarkia*) can be housed in fresh water, at normal room temperature, in small (10-30 gallon) home aquarium tanks. If desired, they have robust reproduction in captivity with scores to hundreds of offspring born from a single egg clutch.

A limited number of papers have shown that crayfish can be used to assess the effects of ethanol on neurotransmission (Swierzbinski et al., 2017) and behavioral effects of other abused drugs, including locomotor responses to intravenous cocaine, morphine and methamphetamine (Imeh-Nathaniel et al., 2017) and the intravenous self-administration of amphetamine (Datta et al., 2018). Furthermore, crayfish are a popular electrophysiology model for introductory neuroscience laboratory classes (Cooper et al., 2011; Ewing and Medler, 2020; Land et al., 2001). Acute serotonin, but not dopamine, injection causes an anxiety-like response in crayfish using a light/dark plus-maze-like assay (Fossat et al., 2015), indicating potential utility as a model of affective disorders.

This study was designed to determine if the crayfish is a suitable model for behavioral studies of the effects of ethanol. Swierzbinski and colleagues showed that if juvenile crayfish are immersed in ethanol (1 M concentration) for 20-40 minutes they exhibit significant behavioral changes, although the behavior was not well described in that paper and an attached movie insinuated pathological levels of intoxication (Swierzbinski et al., 2017). The goal of this investigation was to determine if ethanol would alter locomotor behavior in juvenile crayfish in a dose dependent manner, using immersion time and concentration to alter the dose. The rationale for this study was to use an open field arena to investigate locomotor behavior, with a logic similar to that of the open field in the rodent. In short, crayfish are prey species and have a tendency to avoid staying in open water for long. Crayfish will use tunnels or huts provided in the home aquaria for extended periods of time and will defend them against co-housed conspecifics, if present. Light/Dark transfer models in rodents typically involve light and dark painted or illuminated regions of an arena with open tops in both cases for observation. Pilot studies attempting to adjust the luminance by placing light or dark material under the transparent bins were unsuccessful at generating differential behavior, thus the more naturalistic full-cover was adopted. While this limited full assessment of behavior (e.g., distance traveled, speed, time immobile/mobile) to the open portion, it permitted assessment of time spent under the cover and any transition events (entries), including those between the open and closed portions.

Ethanol was predicted to alter activity of the crayfish, as it does in other laboratory animal models. Bättig first described the effects of ethanol on open field behavior in the rat (Battig, 1969) and subsequent work showed that 0.4 g/kg, EtOH, i.p., increased ambulatory activity and decreased immobility in an open field in rats (Cappeliez and White, 1981). It was found that 2 g/kg, p.o. increases activity in rats, whereas 4 g/kg was reported to decrease activity (Prunell et al., 1987). Similarly 2 g/kg EtOH, i.p., decreased motor activity in 20, 40 or 60 day old rats (Lamble and Rydberg, 1982) and 0.75 g/kg, i.p. depressed horizonal activity in rats (June et al., 1989). Thus it was predicted that lower exposure levels would be associated with increased activity and higher exposure levels with decreased activity in the crayfish.

Ethanol was also predicted to alter anxiety-like behavior in the crayfish. It has been shown that EtOH (1 g/kg, i.p.) in male rats resulted in decreased latency to emerge from the dark portion on a Light/Dark transfer test, increased distance traveled in, and entries into, the light portion of the arena (Sharko et al., 2016). Ethanol in adolescent rats also increased distance and time spent in the open area of an open field, but decreased time spent in the bright portion of a Light/Dark transfer apparatus (Acevedo et al., 2014). Anxiolytic-like effects of EtOH in the Light/Dark test also have been reported for mice (Belzung et al., 1988; Gilmore et al., 1991). Interestingly, a recent report found that adding the selective serotonin reuptake inhibitor citalopram to the housing water decreased the latency of crayfish to emerge from a sheltered location and increased the amount of time spent exploring a food- or conspecific-associated region of an artificial stream environment (Reisinger et al., 2021).

## 2. Methods

### 2.1. Subjects

Crayfish (*Procambarus clarkii*) used in three experimental groups [Cohort 1 (N=15; 9 female / 6 male), Cohort 2 (N=18; 7 female / 11 male), Cohort 3 (N=11; 5 female / 6 male)] were either hatched in the laboratory or obtained from commercial supplier (Carolina Biological Supply). The parents of laboratory bred individuals were obtained from a local pet store. Animals were hatched into a 10 or 20 gallon communal tank and then eventually separated, once reaching ~5 cm in length, into groups (N=3-5) of similarly sized individuals in separated regions (1/3-1/2 of tank size) of a tank. Tanks were equipped with a thin layer of aquarium gravel, 2-4 huts and hideouts, free floating anacharis plants and a continuously running filter. Animals were fed every third day with a rotating variety of foodstuffs including anacharis, frozen krill, shrimp pellets and fish food flakes. During the course of any repeated-measures studies, nail polish markings on the carapace were used to identify individuals.

### 2.2 Behavioral assessment

Locomotor behavior was measured in aquatic open field arenas (see **Figure 1A**). These consisted of 56 cm L × 43 cm W × 15 cm D clear plastic bins placed on a light surface and filled to a 11.5 cm depth with water. Dark cardboard shrouds extended approximately 35 cm above the walls of each arena on all four sides. Locomotor activity was recorded videographically using webcams (Logitech Model C270) mounted approximately 1 meter above the arena. Analysis of the behavioral sessions was conducted off-line using ANY-maze behavior tracking software (Stoelting). Initially the tracking was configured for Center versus the Periphery, with a 34 cm × 18 cm defined Center area, as schematized in **Figure 1B**.

**Figure 1:**
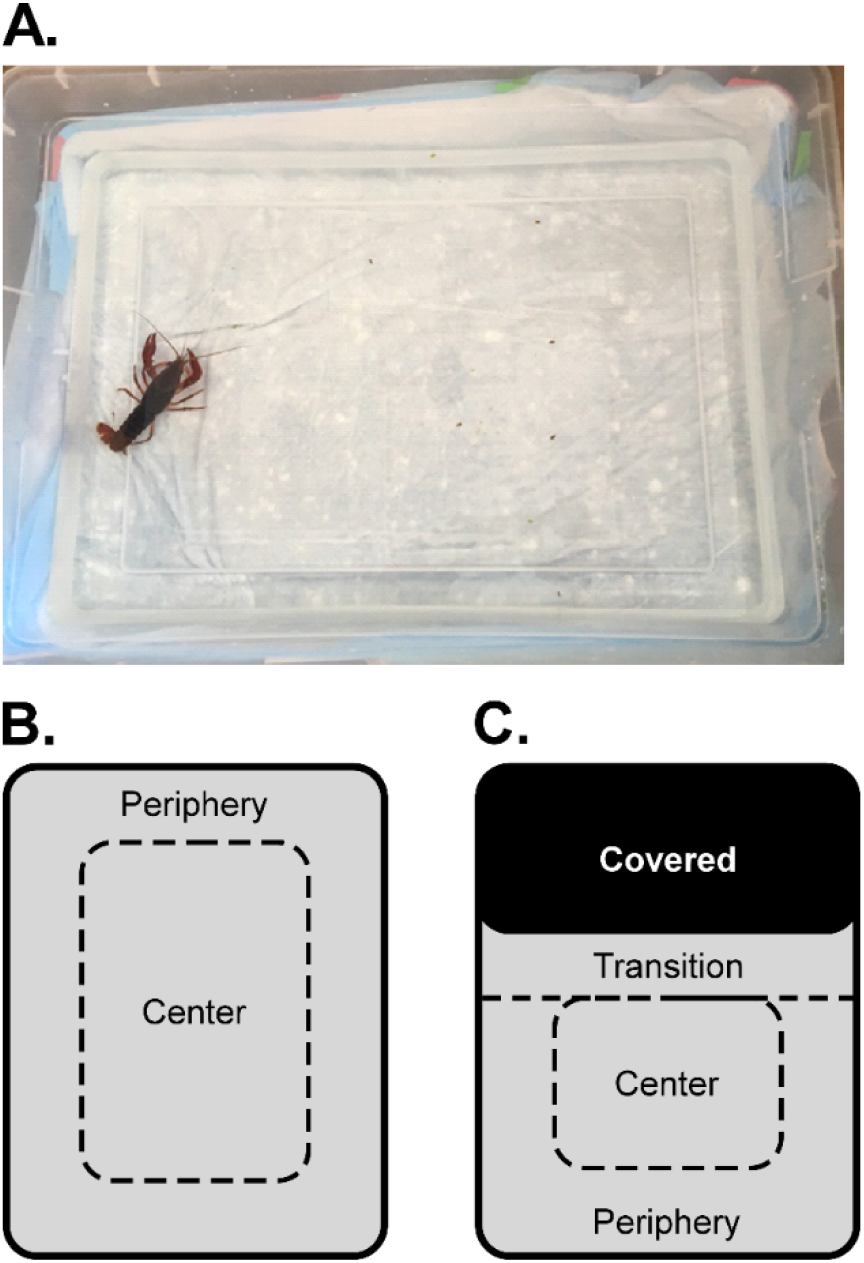
A) A crayfish pictured in the arena. Schematic of B) the open area and C) the arena configured for a light/dark experiment with approximately 1/3 of the area covered. The dotted lines did not appear but approximate where the zones were drawn for the video tracking/scoring.

### 2.3 Ethanol Immersion exposure

Crayfish were exposed to immersion treatment conditions in 16 cm L × 16 cm W × 10 cm H plastic bins filled with 1 L of water by immersion, following the methods described by Swierzbinski and colleagues (Swierzbinski et al., 2017). Immersion conditions included normal aquarium water or aquarium water adulterated with different concentrations of EtOH (0.1, 0.5, 1.0 M) as detailed for specific experiments below. Following immersion, individuals were placed in the Center Zone of the arena and the video recording interval was initiated.

### 2.4 Experiments

#### Experiment 1: Duration of exposure to 1.0 M EtOH

The first study was a baseline recording (no treatment) in the arena for 30 minutes to habituate the animals to the apparatus prior to evaluating the ethanol (EtOH) conditions in a counterbalanced order. For the evaluation of exposure duration, juvenile crayfish (~5-7 cm; ~11 weeks of age) were immersed in water for 20 minutes, or in 1 M EtOH for 10, 20 or 30 minutes and then evaluated in the open field for 30 minutes. The 30-minute maximum interval and the EtOH concentration was selected from a prior paper (Swierzbinski et al., 2017). Treatment conditions were counterbalanced across individuals and conducted no more frequently than every 3-4 days per individual. Two female animals did not complete the study due to mortality during molting, thus N=13 for the analysis.

#### Experiment 2: EtOH concentration

For the evaluation of concentration, the same individual crayfish (N=7 female, N=6 male) were immersed for 30 minutes in water, 0.1 M, 0.5 M or 1M EtOH and then recorded in the open field for 30 minutes. Treatment conditions were counterbalanced across individuals and conducted no more frequently than every 3-4 days per individual. Two of the male animals did not complete the study due to mortality during molting.

#### Experiment 3: Light/Dark Transfer

For the further evaluation of crayfish behavior, one end of the arena was covered ~1/3 with an opaque lid (see **Figure 1C** for schematic). The open area corresponding to Center for the video tracking was now 21.5 cm × 18 cm and there was a new Transition Zone defined, covering an 8.5 cm L rectangle extending from the cover across the width of the arena. This was designed to capture the extension of visible parts of the animal from under the cover when the animal was not fully emerged, based on pilot studies. Those pilot studies also confirmed that the animals do not treat the “edge” defined by the cover across the arena as if it were another wall, i.e., they do not move along it as they do the walls. This video tracking zone mapping also was used to re-analyze locomotor behavior when the arena was completely open for comparison with the behavior when the cover was in place.

Individuals remaining from the same group (N=6; 4 female / 2 male) of crayfish used in Experiment 2, now adult and with prior EtOH and arena experience, were used to pilot the Light/Dark arena approach and to test the hypothesis that crayfish would spend time preferentially under the cover. The cover was placed in a fixed location for all individuals in this study, i.e., at the rear relative to where the experimenter faced the arena and placed/removed the animals.

#### Experiment 4: Effects of Repeated immersion on Light/Dark Transfer

For this experiment adult crayfish (>7 cm) were immersed for 30 minutes each day for 5 sequential days. Locomotor testing in the Light/Dark Transfer configuration was conducted after the first (N=18 EtOH; N=11 Water), third (N=12 EtOH; N=11 Water) and fifth (N=16 EtOH; N=11 Water) days. Locomotor testing was also evaluated on Day 8 after 30 minutes of water immersion (all individuals).

#### Experiment 5: Hemolymph EtOH concentration

Hemolymph samples were collected from the pericardial sinus of adult crayfish in EDTA tubes following exposure to 0.1 and 1.0 M EtOH for 30 minutes. Samples were centrifuged for 10 minutes at 5,000 G and the supernatant was collected, frozen (−80 °C), and stored for later assessment. Ethanol concentrations were thereafter determined by GC-MS (Agilent 7820A GC coupled to a 7697A headspace sampler, Agilent Technologies, Santa Clara, CA, USA). Crayfish were euthanized immediately after hemolymph sample collections.

### 2.5 Data analysis

ANY-maze software was used to score behavior in the Center versus the surrounding Periphery zone for the Open Field Arena (Figure 1B), and in the Center, Periphery, Covered and Transition zones for the Light/Dark Arena (Figure 1C). For the Open Field arena assessments, the dependent measures included Distance Traveled, Percent of the Time spent Immobile, Speed, Center Entries, and Time spent in Center versus Periphery of the arena. Treatment of missing data (e.g., due to an animal failing to enter the peripheral zone at all, failing to ever stop moving in a zone, or being unavailable for a given treatment condition) was handled by using mixed-effects analysis. One-way ANOVA (or mixed-effects) was used for overall Distance Traveled, Center Entries and Center Time with the single factor of Immersion Condition. Two-way ANOVA (or mixed-effects) was used with factors of Zone and Immersion Condition to analyze Distance Traveled by zone, Speed and Percent of the Time spent Immobile. Initial analysis considered the entire sample and limited follow-up analysis divided the sample into male and female individuals to assess any possible sex differences. For the Light/Dark arena validation, the dependent measures include Distance Traveled, Speed of travel, Percent Time spent, and Time spent Immobile, by zone.

In the repeated immersion study, the dependent measures for the Light/Dark arena included Distance Traveled, Speed of travel, Time spent in a given zone and the number of Entries into a given zone. The first analysis contrasted the treatment groups on Day 1 and Day 8, thus there was a repeated measures factor of Zone and a between-subjects factor of Immersion condition (Water vs. EtOH). The data then were analyzed within-group by treatment Day (D1-D8) because the first hypothesis under investigation involved tolerance with repeated administration (D1 to D5 as the critical test) and any effect of discontinuation (i.e., D8 compared with D1 and D5). A D3 test was included for most of the study because the initial cohort (N=6) exhibited no difference between D1 and D5 and thus there was a concern a biphasic trend was being missed. Thus, the design included repeated measures factors for Zone and assessment Day, and again, a mixed-effects analysis was used to account for missing subjects in a given treatment condition or behavioral measure. Any significant effects were followed with post-hoc analysis using Tukey correction for all multi-level, and Sidak correction for any two-level, comparisons. Prism 9 for Windows (v. 9.1.1; GraphPad Software, Inc, San Diego CA) was used for all analyses.

## 3. Results

### 3.1 Baseline behavior

The *Procambarus clarkii* juvenile crayfish exhibit significant locomotor behavior under baseline conditions when placed in the brightly illuminated open field, traveling an average of 23 (SEM 2.9) meters in 60 minutes (**Figure 2A**). The distribution of the animals’ time and activity in the Center versus the Periphery under baseline conditions showed a distinct preference for the Periphery. They moved faster (2-3 x) when in the center (**Figure 3A**) while spending less time (~90 out of the 1800 second session; **Figure 3D**) and moving less (~10%) total distance (**Figure 2B, C**) in the Center Zone. These relative patterns were consistent across all treatment conditions in the EtOH experiments, as well.

**Figure 2:**
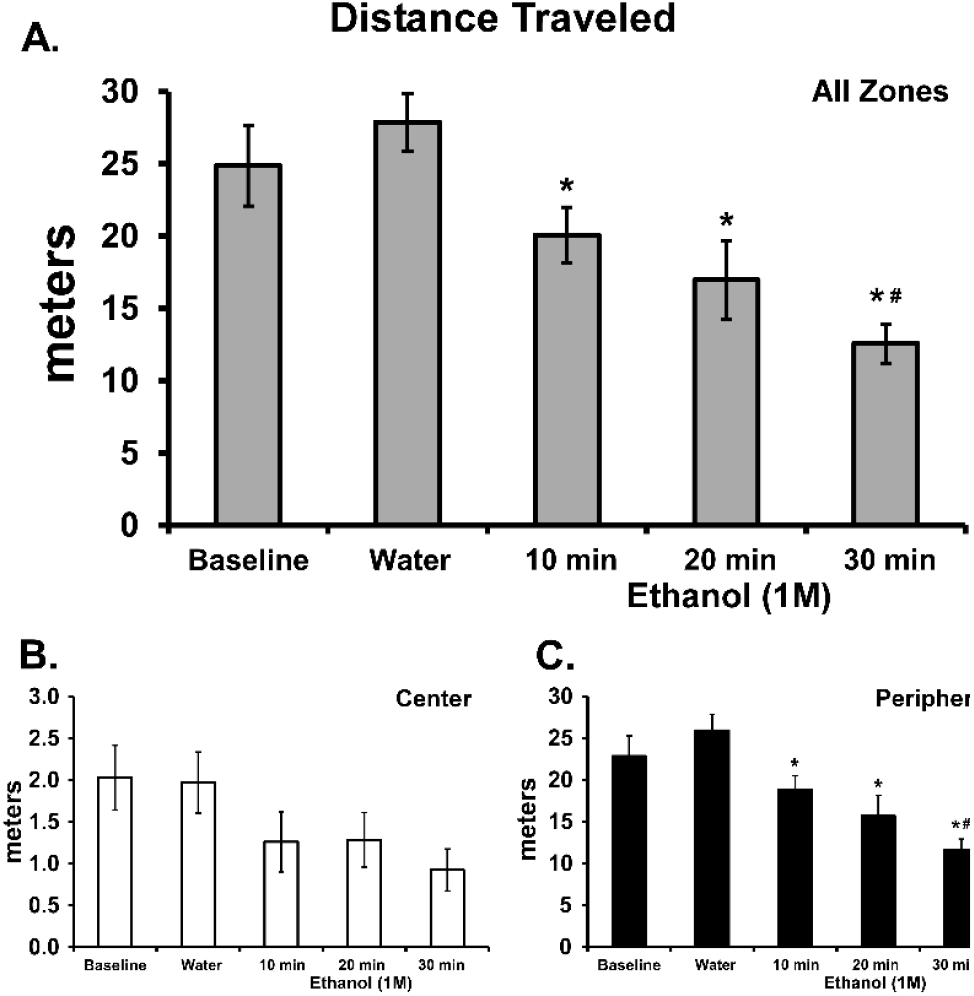
Mean (N=13; ±SEM) distance traveled by crayfish in the open arena over 30 minutes. N.b. There is a 10-fold difference in the scale for Center and Periphery. A significant difference from Water is indicated with * and a difference from 10 minutes with #.

**Figure 3:**
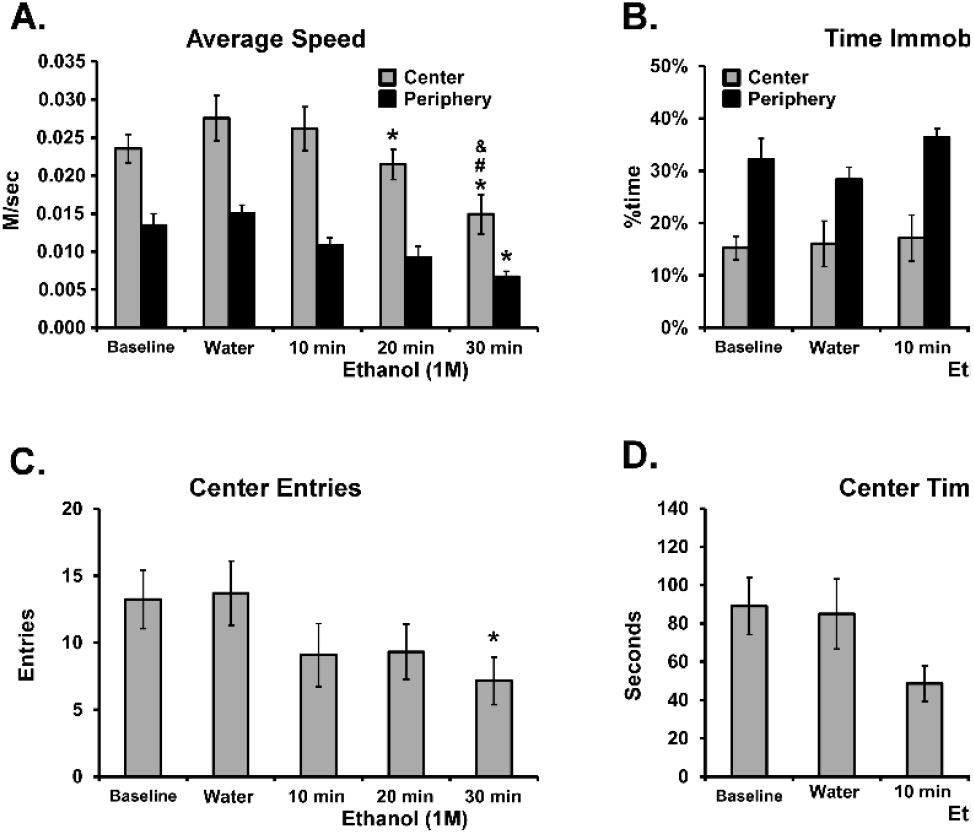
Mean (N=13; ±SEM) speed, time spent immobile, center entries and time spent on the center of the open arena over 30 minutes. A significant difference from Water is indicated with * and a difference from 10 minutes with # and a difference from 20 minutes with &.

### 3.2 Effects of Ethanol Exposure Duration

The effects of ethanol on locomotor behavior were immersion-interval dependent in a monotonic relationship for most activity measures. Following immersion in EtOH (1.0 M) for 30 minutes, crayfish traveled approximately half of the distance traveled after water immersion, with an approximately similar magnitude of reduction in both Center and Peripheral zones (**Figure 2**). The statistical analysis confirmed significant effects of EtOH immersion on Distance Traveled overall [F (2.817, 33.80) = 13.40; P<0.0001; **Figure 2A**] and Center Entries [F (2.769, 33.23) = 4.84; P<0.01; **Figure 3C**]. Crayfish <u>*traveled significantly more distance*</u> in the Periphery, compared with the Center, as confirmed in the analysis [significant effects of Zone, F (1, 12) = 199.9; P<0.0001; Immersion Condition, F (4, 48) = 13.42; P<0.0001; and the interaction, F (4, 48) = 11.68; P<0.0001; **Figure 2 B, C**]. The post-hoc test confirmed significant reductions in peripheral distance traveled, relative to water immersion, for all three EtOH durations and also after 30 minutes compared with 10 minutes of EtOH immersion. In addition, significantly less distance was traveled in the Center versus the Periphery after each of the immersion conditions. The crayfish <u>*moved faster*</u> in the Center compared with the Periphery [significant effects of Zone, F (1, 12) = 107.7; P<0.0001; Immersion Condition, F (4, 48) = 8.93; P<0.0001; interaction, n.s.; **Figure 3 A**]. The post-hoc test confirmed that movement was faster in the Center following each of the Immersion conditions, and that significantly slower movement was observed in the Center after 20 or 30-minute immersion, and in the Periphery after 30 minutes of immersion, relative to water immersion. Slower movement in the Center was also confirmed after 30 minutes of immersion compared with 10- or 20 minutes. The crayfish spent a larger proportion of their <u>*time immobile*</u> when in the Periphery compared to when in the Center (significant effects of Zone, F (1, 12) = 48.97; P<0.0001; Immersion Condition, F (4, 48) = 5.09; P<0.005; interaction, n.s.; **Figure 3 B**). The post-hoc test confirmed that less time was spent immobile in the Center across all Immersion conditions, and that significantly more time was spent immobile in the Center after the 30-minute immersion, and in the Periphery after 20 or 30 minutes of immersion, relative to water immersion.

### 3.3 Effects of Ethanol Concentration

In the second experiment, the effects of ethanol depended on the concentration, again in a monotonic relationship for most activity measures, including overall Distance Traveled [F (2.335, 23.35) = 10.65; P<0.0005; **Figure 4**]. There was no significant effect of immersion condition on time spent in the Center (**Figure 5 D**) or Center entries (P<0.06; **Figure 5 C**). Crayfish <u>*traveled significantly more distance*</u> in the Periphery, compared with the Center, of the arena and this distance was altered by EtOH immersion [significant effects of Zone, F (1, 10) = 96.04; P<0.0001; Immersion Condition, F (3, 30) = 10.64; P<0.0001; and the interaction, F (3, 30) = 9.98; P=0.0001; **Figure 4 B, C**]. The post-hoc test confirmed a significant reduction in peripheral distance traveled after 1.0 M EtOH immersion, relative to all other immersion conditions. There was also a significant difference in between the 0.1 M and 0.5 M EtOH immersion conditions. In addition, the post-hoc test confirmed that significantly less distance was traveled in the Center versus the Periphery for each of the immersion conditions.

**Figure 4:**
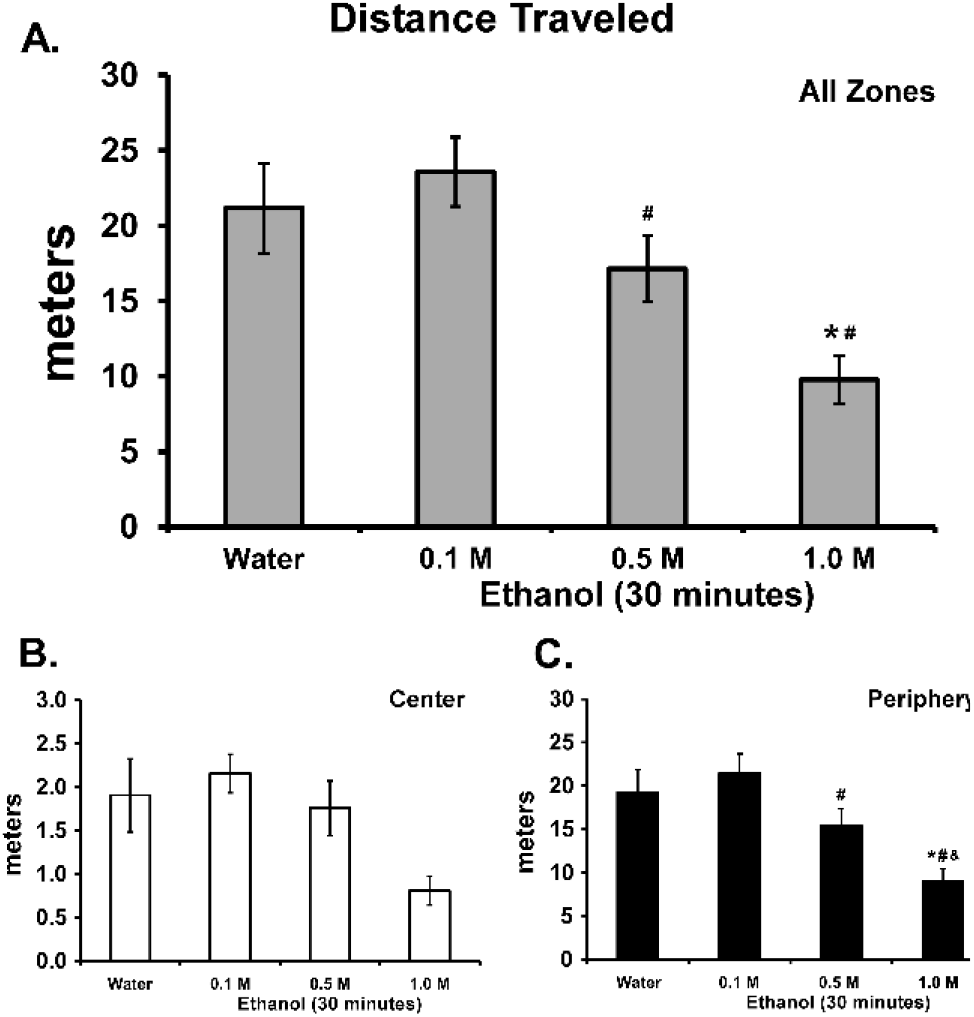
Mean (N=11; ±SEM) distance traveled by crayfish in A) all of the open arena, or in the B) center and C) peripheral zones, over 30 minutes. N.b. There is a 10-fold difference in the scale for Center and Periphery. A significant difference from Water is indicated with *, a difference from 0.1 M immersion with # and from 0.5 immersion with &.

**Figure 5:**
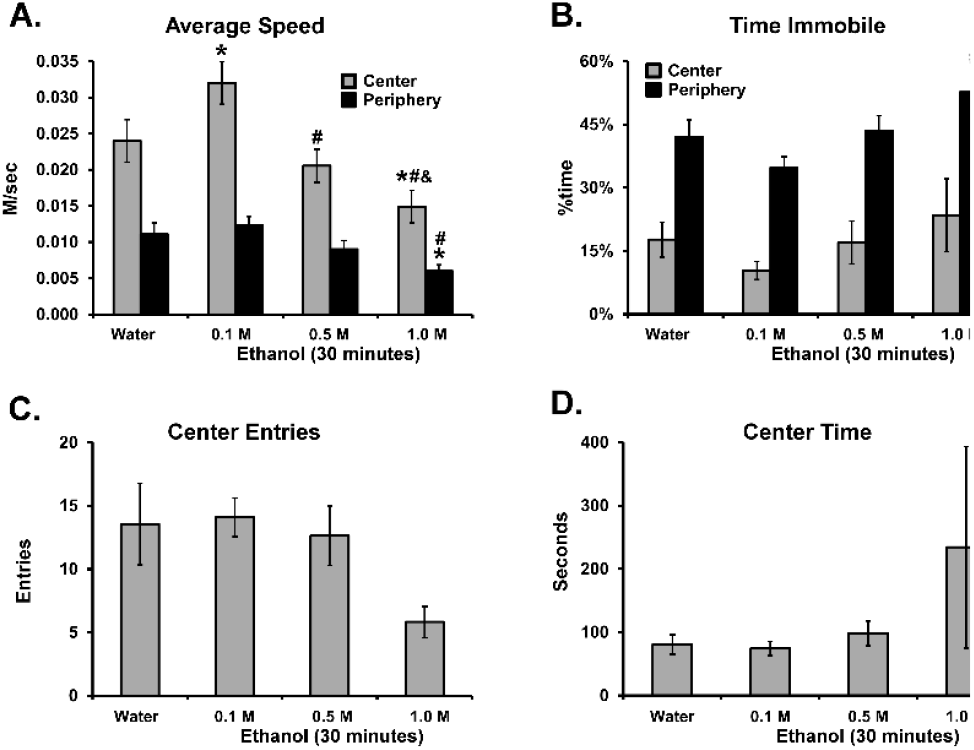
Mean (N=11; ±SEM) A) speed of travel, B) time spent immobile, C) entries into the center zone and D) time spent in the center zone, in the open arena over 30 minutes. A significant difference from Water is indicated with *, a difference from 0.1 M immersion with # and a difference from 0.1 M immersion with &.

The crayfish <u>*moved faster*</u> in the Center compared with the Periphery [significant effects of Zone, F (1, 10) = 104.2; P<0.0001; Immersion Condition, F (3, 30) = 11.44; P<0.0001; and of the interaction, F (3, 30) = 5.30; P<0.005; **Figure 5 A**]. The post-hoc test confirmed that movement was faster in the Center for each of the Immersion conditions, and that significantly slower movement was observed in the Center after 1.0 M EtOH immersion compared with all other conditions. Movement in the Center was also faster after the 0.1 M EtOH immersion compared with all other immersion conditions. Movement speed was slower in the Periphery after 1.0 M EtOH immersion compared with water or 0.1 M EtOH immersion. The crayfish spent a larger proportion of their <u>*time immobile*</u> when in the Periphery compared to when in the Center (significant effects of Zone, F (1, 10) = 51.08; P<0.0001; Immersion Condition, F (3, 30) = 3.289; P<0.05; interaction, n.s.; **Figure 3B**). The post-hoc test confirmed that less time was spent immobile in the Center after each of the Immersion conditions, and that significantly more time was spent immobile in the Periphery after 1.0 M EtOH immersion compared with 0.1 M EtOH immersion.

### 3.4 Hemolymph Ethanol Levels

Hemolymph was collected from crayfish following 30-minute immersion in EtOH 0.1 M (N=3) and 1.0 M (N=4) baths. Two additional untreated crayfish samples were obtained as controls (and had 0 EtOH). The blood hemolymph levels (EtOH mg/dL) are depicted in **Figure 6** and show a concentration dependent increase in circulating EtOH (Unpaired t-test: t=6.273, df=5, P<0.005).

**Figure 6:**
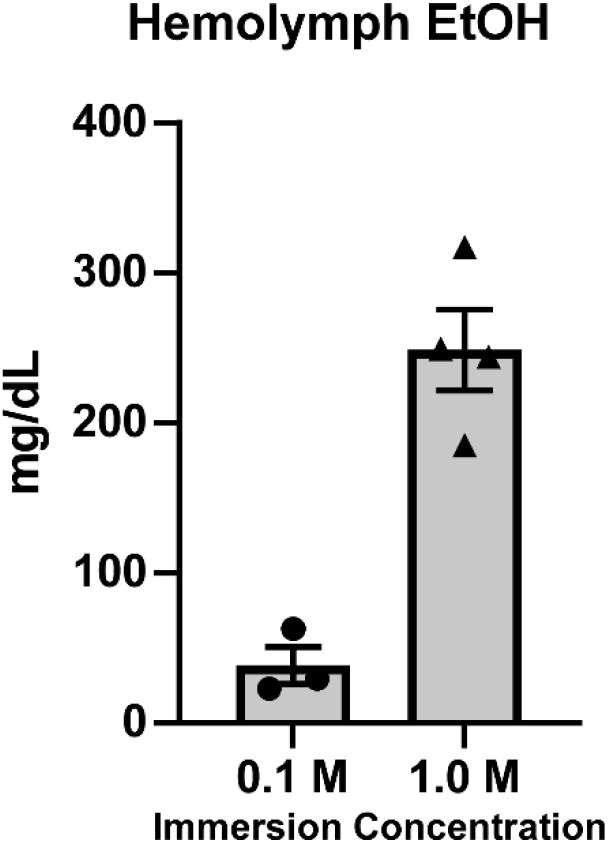
Mean (±SEM) concentration of EtOH in the hemolymph of crayfish following 30-minute immersion in 0.1 M (N=3) or 1.0 M (N=4) EtOH.

### 3.5 Light/Dark Transfer Arena

Under baseline (no immersion) conditions the remaining adult crayfish (N=6; 2 male) spent the majority of their time (mean 67%; range 43%-85%) under the cover in the Light/Dark arena. Their behavior was significantly affected by Zone when in the uncovered regions, as the analysis confirmed significant effects on Distance Traveled (F (1.016, 5.078) = 14.40; P<0.05; **Figure 7 A**) and Speed (F (1.306, 6.532) = 13.98; P<0.01; **Figure 7 B**), but not Time spent Immobile (P=0.74; **Figure 7 C**). The post-hoc tests confirmed that they traveled the most distance in the Periphery (**Figure 7 A**) and moved significantly faster in the Center Zone compared with Periphery or Transition Zones (**Figure 7 B**). The patterns of behavior in the open part of the arena were therefore similar to the patterns when the arena was entirely uncovered.

**Figure 7:**
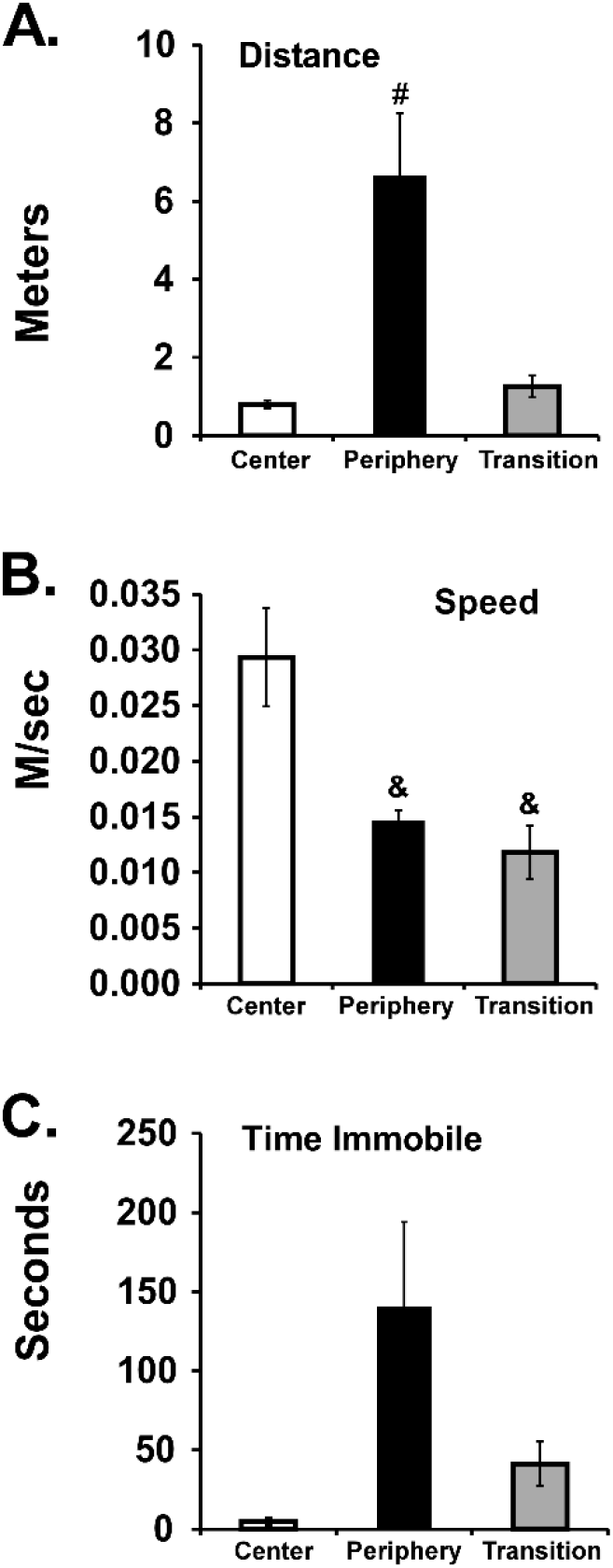
Mean (N=6; 4 female; ±SEM). Mean distance traveled, B) speed of locomotion and C) Time immobile across the zones. A significant difference from the center zone is indicated with &, and from all other zones with #.

An additional analysis was done to compare the percent time spent under the cover when placed over the back third for this Light/Dark configuration, compared with the distribution of time when in the Open Field arena configuration. The open arena baseline recordings (from Experiment 1) for the sample of 6 animals were re-scored with the ANY-maze template used for the covered run. These videos were first scored with the virtual “cover” at the rear and then were scored again with the location of the virtual cover switched to the front. The first important observation is that crayfish spent differing amounts of time in different zones (**Figure 8 A**), as confirmed with a main effect of Zone [F (3, 15) = 154.0; P<0.0001] on the percent of time spent. The post-hoc test confirmed that Percent Time differed between all zones except Center and Transition when the cover was in place. An identical result was confirmed for the virtual cover re-scoring when the cover was in the rear. A similar result was also confirmed for when the virtual cover was in the front, except that Percent Time in Periphery did not differ from Percent Time under the (virtual) Cover. Importantly, the animals spent more time in the Covered Zone *when there was an actual cover in place*. This was confirmed with a significant interaction of Zone with Scoring template [F (6, 30) = 12.24; P<0.0001]. The post hoc test further confirmed that Percent Time in the Periphery and in the Covered portion of the area differed significantly between the actual Light/Dark condition and each of the virtual cover analysis conditions. Thus, the cover caused a significant increase in the amount of time spent in that part of the arena. There was nothing unusual about the six animals used for the Light/Dark pilot studies in terms of distribution of behavior in the open arena. Using the full sample (N=14; 6 male) which had been run in the baseline condition in the first Open Arena experiment and re-scoring the data with both front and back virtual covers, the same pattern of time distribution was confirmed (**Figure 8 B, C**). Moreover, there was no difference between the sexes and no consistent preference for the front or rear of the arena (again, the cover is a virtual scoring template and did not exist on the arena during this recording session). Therefore, the hypothesis that crayfish would preferentially avoid the open areas when provided with a cover was supported.

**Figure 8:**
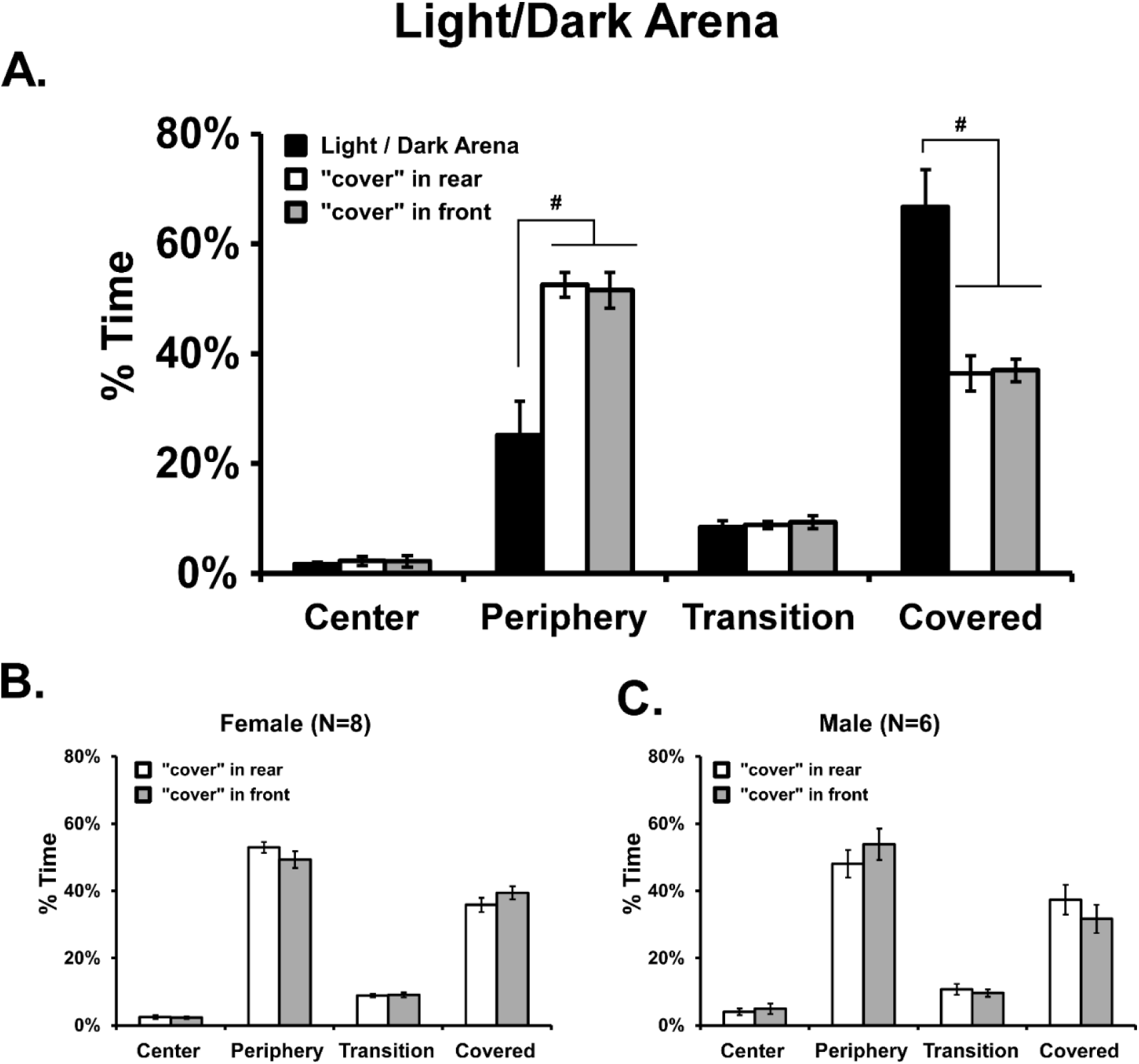
A) Mean (N=6; 4 female; ±SEM) percent of time spent in four zones of the Light/Dark arena and in the corresponding areas of the Open Arena during baseline assessment. B) Mean percent of time spent in areas of the Open Arena corresponding to Light/Dark zones for B) female (N=8; ±SEM) and C) male (N=6; ±SEM) crayfish during baseline assessment. A significant difference between scoring runs is indicated with #.

### 3.6 Effects of Repeated EtOH on Light/Dark Arena Activity

For the group (N=18, 7 female, for Day 1, N=12, 4 female, on Day 3 and N=16, 3 female, on Days 5 and 8) of crayfish treated with five sequential days of EtOH (1.0 M) immersion for 30 minutes and then assessed after water immersion three days later, the EtOH significantly altered behavior in the Light/Dark arena (**Figure 9**). First, the analysis of activity on Day 1 with the Water (N=11, 5F) immersion group confirmed that there was a significant effect of Zone (F (1.327, 47.78) = 130.6, P<0.0001) and of the interaction of Zone with Immersion condition (F (3, 108) = 8.534, P<0.0001) on Time spent. The post-hoc test confirmed that the EtOH animals spent less time under the Cover and more time in the Periphery Zone compared with the Water immersion group. No group differences were confirmed for Entries, Speed or Distance on Day 1.

**Figure 9:**
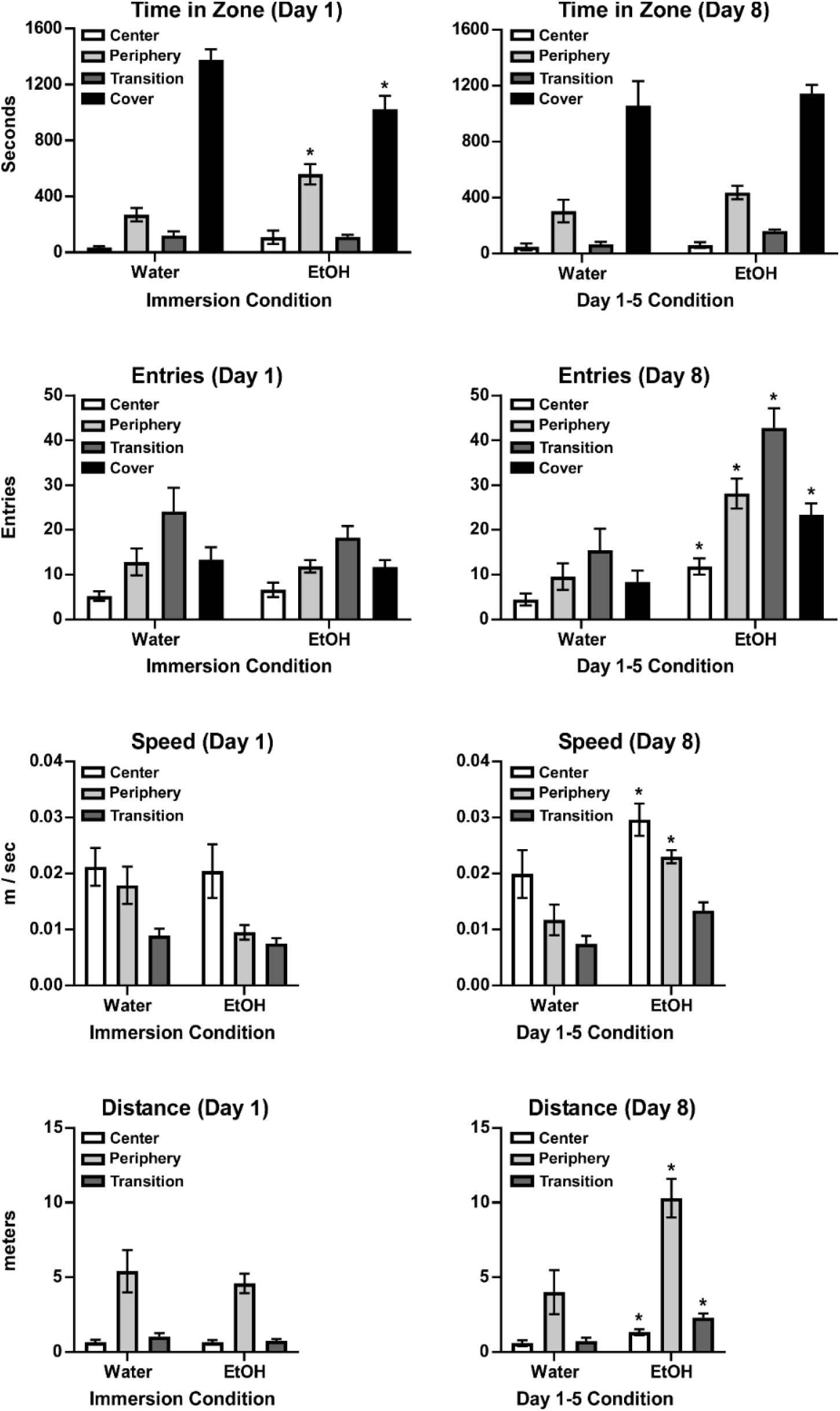
Mean (±SEM) time spent, entries, speed of travel and distance traveled by zone for groups immersed in Water [N=11 (5 female] or EtOH [N=18 (7 female) on Day 1, N=16 (6 female) on Day 8] for 5 days. A significant difference between immersion condition is indicated with *.

On Day 8, however, there was a significant effect of Zone (F (1.384, 46.15) = 102.7; P<0.0001), but not any significant effect of the interaction of immersion group with zone on Time spent (P=0.09). However, there were significant effects of immersion Group, and/or the interaction of Group with Zone on Entries [Group, F (1, 25) = 16.73, P<0.0005; Zone, F (1.403, 35.07) = 53.92, P<0.0001; Interaction, F (3, 75) = 12.26, P<0.0001], Speed [Group, F (1, 25) = 8.390, P<0.01; Zone, F (1.596, 39.10) = 41.45, P<0.0001; Interaction, *n.s.*] and Distance [Group, F (1, 25) = 16.73, P<0.005; Zone, F (1.403, 35.07) = 53.92, P<0.0001; Interaction, F (3, 75) = 12.26, P<0.001] on Day 8. The post-hoc test confirmed significant differences between immersion groups on Entries (all zones), Speed (Center and Periphery) and Distance (all zones). The within-group analyses across the four recording Days in the Water immersion group did not confirm any significant effects of Day, nor any interaction of Day with Zone, for Time spent, Entries, Speed or Distance traveled (**Figure 10**).

**Figure 10:**
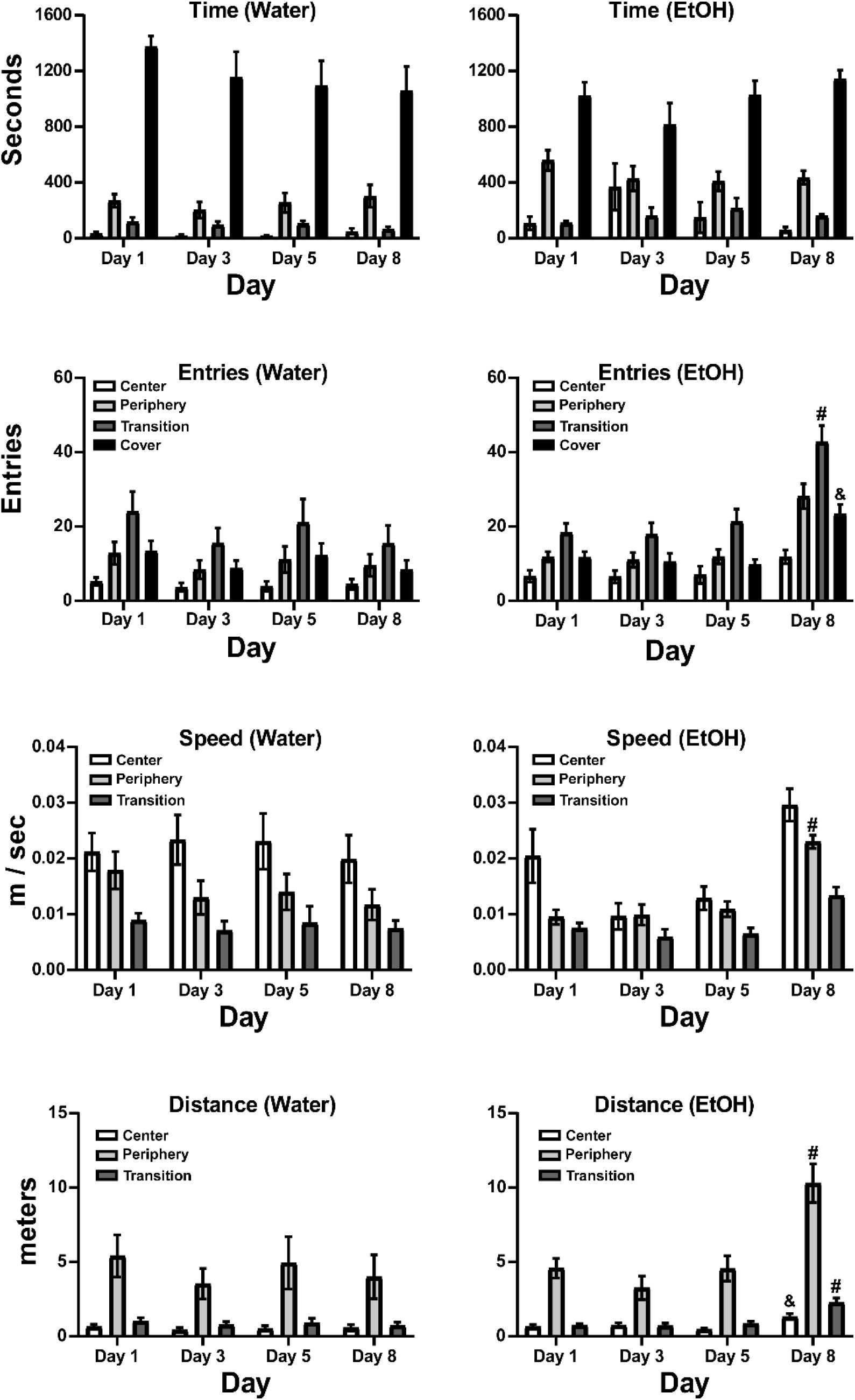
Mean (±SEM) time spent, entries, speed of travel and distance traveled by zone and Day for groups immersed in (left panels) Water (N=11, 5 female) or (right panels) EtOH (N=18, 7 female on Day 1, N=16, 6 female on Day 8) for 5 days. A significant difference from all other days, within Zone, is indicated with #, and from Day 5, within Zone, with &.

In the EtOH immersion group, there were no significant effects of Day, nor of the interaction of Day with Zone, for Time spent. However, there were significant effects confirmed for Entries [Zone: F (1.885, 32.05) = 45.09, P<0.0001; Day: F (2.246, 38.18) = 15.76, P<0.0001; Interaction: F (3.611, 43.74) = 6.741, P<0.0005], Speed [Zone: F (1.139, 19.36) = 16.75, P<0.0005; Day: F (2.087, 35.48) = 14.14, P<0.0001; Interaction: n.s.] and Distance traveled [Zone: F (1.037, 17.63) = 59.51, P<0.0001; Day: F (2.127, 36.16) = 18.14, P<0.0001; Interaction: F (1.876, 21.89) = 11.04, P<0.001]. As depicted on **Figure 10**, the post-hoc tests confirmed that these effects were due to increased entries, speed and distance traveled on Day 8 (after water immersion) as compared with prior days in which EtOH immersion was conducted prior to locomotor assessment.

## 4. Discussion

The experiments show that the Red Swamp Crayfish, *Procambarus clarkii*, is a suitable model for the evaluation of behavioral effects of ethanol (EtOH). Administration by bath immersion results in levels of EtOH in the hemolymph that approximate blood EtOH concentrations that are significant in laboratory mammals as well as in humans. Dose-dependent effects can be produced by altering the concentration of ethanol in the immersion bath or by changing the amount of time in which they are immersed in a given concentration. The EtOH immersion was shown to alter locomotor behavior in an aquatic version of an open field arena with locomotor suppression produced in a dose dependent manner. There was also some evidence of a locomotor stimulant effect at the very lowest dose. Crayfish exhibited zone preference in both the Open Field and in the partially covered, Light/Dark version of the arena in a manner consistent with avoidance of the open parts, similar to preferences expressed by laboratory rodents on similar assays. Finally, immersion in EtOH produced an anxiolytic-like effect in the Light/Dark by decreasing time spent under the cover.

### 4.1 Open Field

Crayfish behavior in the Open field was differentially distributed by zone in a manner that avoided the Center area. That is, they moved more quickly in the Center Zone and traveled about 1/10 of the distance within the Center, compared with the Peripheral Zone. They also spent approximately half as much of their time immobile in the Center, as compared with the Periphery. There was no clear evidence of an anxiolytic or anxiogenic effect of EtOH immersion in the Open Field experiment, since there were no consistent shifts to greater preference for, or avoidance of, the Center produced. While fewer entries into the Center were made in the 30-minute immersion / 1.0 M EtOH condition, the animals also spent more time immobile in the Periphery. In addition, there was a monotonic dose-related decrease in distance traveled in both Center and Peripheral zones and a decrease in the speed of movement as well. This suggests that the reduction in Center entries associated with EtOH immersion is more likely attributed to the general reduction in locomotor activity, and not with altered sensitivity to the open space. There were no sex differences observed as outlined in the Supplemental Materials (**Supplemental Figures S1-S4**). There was evidence of a small locomotor stimulant effect of 30-minute immersion in 0.1 M EtOH as reflected in a significant increase in Center speed. This is consistent with observations in the rat in which lower doses produce locomotor stimulation while higher doses suppress activity (Prunell et al., 1987). There was no indication of a more robust stimulant effect of 0.05 M immersion, see **Supplemental Figure S5**.

### 4.2 Hemolymph Ethanol Concentration

The hemolymph EtOH concentrations averaged 38.4 mg/dL (0.038 HAC) after immersion in 0.1 M EtOH and 248.9 mg/dL (0.249 HAC) after immersion in 1.0 M EtOH. This corresponds well with relevant blood EtOH levels in vertebrate animals, including mice, rats (de Guglielmo et al., 2017; Dilley et al., 2018; Ginsburg et al., 2008), monkeys (Crean et al., 2011; Katner et al., 2007; Wright and Taffe, 2014) and humans (Irwin et al., 2013; Phillips et al., 2015), after exposure to EtOH by the oral route of administration. The lower end of this range is associated with a threshold for observable behavioral effects and the high end is associated with significant intoxication. Importantly, this is the range of blood EtOH concentrations reached in *voluntary*, self-administration approaches in rat and monkey models (de Guglielmo et al., 2017; Katner et al., 2004; Wright and Taffe, 2014).

### 4.3 Light/Dark Test

The Light/Dark transfer test results further indicated that crayfish avoid the open and indicated that they prefer the dark or covered portion of an environment, when it is available. In the initial experiment, crayfish spent two-thirds of their time in approximately one-third of the arena (by area or by peripheral wall length) under baseline, or untreated conditions. In contrast, they spent 36% of their time in the same location when the arena was entirely open. Thus, the study found that the distribution of zone time in the Open arena (Center versus Periphery) and in the Light/Dark arena (Covered versus Center or Periphery) may serve as an anxiety-like measure akin to the analogous tests used in rodents. The utility of the assay was further supported by the lack of change in the pattern of behavior across the four sessions for the Water control group in the repeated-immersion experiment. That is, time spent in each zone, zone entries, speed and distance traveled did not change over the four test sessions. There was, however, a consistent effect of zone across all three measures and, again, animals spent most of their time under the cover and the least amount of time in the center zone. Their speed was fastest in the center zone and slowest in the transition zone, while they traveled the most distance in the periphery. The fewest entries were made in the center zone as well. Importantly, this confirms the behavioral stability over repeated testing within a single group.

### 4.4 Effect of Repeated EtOH on Light/Dark Behavior

There was evidence for an anxiolytic effect of EtOH immersion in the Light/Dark experiment. The EtOH group spent significantly less time under the Cover and more time in the Peripheral zone compared with the Water control group on Day 1. Zone Entries, Speed and Distance Traveled did not differ significantly between the groups on Day 1, ruling out explanations based on general locomotor suppression.

The crayfish did not develop tolerance (or sensitization) to EtOH when the group was exposed daily for five days, as no substantial change in behavior was observed between Day 1 and Day 5 of the repeated dosing experiment. More specifically, the pattern of fewer entries into the Center Zone, more entries into the Peripheral and Covered Zones and the most entries in the Transition Zone persisted across the EtOH (Day 1, 3 and 5) sessions with no difference across the days. Likewise, crayfish traveled the most distance in the Peripheral zone and the least in the Center zone across all observation days. One potential difference associated with repeated EtOH immersion was the observation that Center speed was not higher than the speed of travel in the other zones on Day 3 and Day 5, as it was on the water day and in general across all the experiments reported herein. There were no other similar consistent differences that emerged only on Day 3 and Day 5, making this difficult to interpret other than it may reflect an anxiolytic effect of EtOH which was obscured on the Day 1 due to the novelty of the procedure. Although two outliers (0.06 and 0.082 m/sec respectively) contributed to the high Center speed average on Day 1 for the EtOH group, removal of these two values did not eliminate the pattern of increased Center speed. Results of analyses either between groups (**Figure 9**) or within the EtOH treated group (**Figure 10**) revealed increased zone entries, speed and distance traveled after three days of abstinence from repeated EtOH immersion (Day 8). Despite the repeated EtOH having no substantial differential effect on Zone Entries, Speed and Distance Traveled compared with the water group on Day 1, there was an excess of activity in the repeated EtOH group compared with the water group on Day 8, which was specific to the discontinuation from repeated EtOH. Although the neuropharmacological underpinnings of this phenomenon are as yet unknown, this highlights the utility of this model to examine short (and possibly long) term effects of abstinence from repeated EtOH exposure.

### 4.5 Replication and Reliability

Although the study constitutes an initial development effort, it had a set of design features that address replicability and stability of the observed effects. First, the Open Field experiment generated similar results across the baseline (first arena exposure) and the water immersion condition (which was conducted in a randomized order with the EtOH immersion in the time-of-immersion study). Second, the dose-immersion replication of the 1M 30-minute immersion across the two Open Field experiments produced a similar magnitude of effect. Finally, no significant sex differences were observed in either baseline behavior, i.e., in the effects of ethanol on locomotor behavior or in the distribution of behavior in the virtual cover re-scoring of the baseline activity. This constitutes an important sub-group replication.

This study was not designed to provide developmental information on the locomotor effects of ethanol immersion, given the sequential study in the same animals and no attempt to define developmental age beyond length and approximate age. Nevertheless, the effect of 1.0 M EtOH immersion for 30 minutes was nearly identical across the first two experiments when animals had grown from late juvenile to early adult length.

### 4.6 Advantages of Crayfish Models

Crayfish species are widely distributed, worldwide, in natural environments (https://en.wikipedia.org/wiki/Crayfish). Subjects for use in laboratory experiments can be purchased from pet stores (as we did for some of our subjects), biological suppliers (Swierzbinski and Herberholz, 2018; Teshiba et al., 2001), or are easily fished from the wild, if locally available (Datta et al., 2018; Imeh-Nathaniel et al., 2017). This invertebrate species can be used for research within institutions that do not maintain regulatory approvals and facilities appropriate for vertebrate animal research subjects. The marbled crayfish, *Procambarus virginalis*, (“Marmokrebs”) is a parthogenic species; since individuals are genetic clones, and the *P. virginalis* genome was sequenced in 2018 (Gutekunst et al., 2018) and genetic experiments might be facilitated. The fact that these species have external egg incubation might be leveraged with Cas9-mediated gene editing techniques, e.g., (Chaverra-Rodriguez et al., 2018). In addition, crayfish are a popular electrophysiology model for introductory neuroscience laboratory classes (Cooper et al., 2011; Ewing and Medler, 2020; Land et al., 2001), and the present studies potentially expand the scope of behavioral and pharmacological experiments that could be adapted. More simply, crayfish are often maintained as classroom pets in primary schools in the US. This latter familiarity in the educational environment may support demonstration labs, and/or science fair projects in primary and secondary educational settings.

### 4.7 Alternative Approaches

The materials necessary for these experiments are relatively inexpensive and are, for the most part, available from local pet stores, home goods, liquor / grocery and electronic stores. The most expensive component of these studies is the video tracking suite, ANY-maze, which could potentially be replaced with open source software such as ToxTrac (Rodriguez, A., Zhang, H., Klaminder, J., Brodin, T., Andersson, P. L. and Andersson, M. (2018). ToxTrac: a fast and robust software for tracking organisms. Methods in Ecology and Evolution. 9(3):460–464; https://sourceforge.net/projects/toxtrac/) or DeepLabCut (http://www.mackenziemathislab.org/deeplabcut).

## Acknowledgements

The authors would like to acknowledge the efforts of Mitchell L. Turner and Rachelle N. Tran who conducted many preliminary experiments to develop and refine the methods for evaluating the behavioral effects of drugs in crayfish. We are likewise grateful to Michael Userenko for assistance with conducting some experiments and to COJT for donation of nail polish to identify animals. We are grateful to Professor Zen Faulkes, Ph.D., for critical initial advice on crustacean handling, immobilization and euthanasia. Hemolymph alcohol assessment was conducted with resources provided by the TSRI Alcohol Research Center (P60 AA006420). Author AG was supported by T32 AA007456 and by a UCSD Chancellor’s Post-doctoral Fellowship.

