## Supplemental Materials for "Behavioral effects of ethanol in the Red Swamp Crayfish (*Procambarus clarkii*)"

***A crayfish model for assessment of the behavioral effects of alcohol***

### 1. Supplemental Introduction

#### Sex Differences:

Consistent with recent calls to increase the consideration of sex as a biological variable that might be relevant to numerous scientific inquiries (McGregor et al., 2016; Shansky and Murphy, 2021), we tracked the sex of the animals used in our studies. For a few key conditions we dis-aggregated by sex to determine any differences in behavior. The analysis of light/dark behavior for the validation study was presented in Figure 8 of the main report. Here we present the measures for the Open Field arena disaggregated by sex.

#### Effects of EtOH on Light/Dark transfer:

The initial validation of the Light/Dark transfer was conducted in experimentally experienced individuals who were also older than the group used for the initial Open Field challenge. In addition, the concentration experiments in the Open Field were conducted in the first group after the duration of exposure experiment. Thus, we conducted two experiments in which experimentally naïve crayfish of the ~5-7cm length were exposed to different EtOH concentrations for 30 minutes prior to the session. The first cohort was exposed to 0.05, 0.1 and 1.0 M EtOH immersion because the concentration experiment in the Open Field found a slight stimulatory effect of 0.1 M EtOH immersion and thus exploration of the lower end was of interest. The results of this experiment, however, did not produce a graded dose-effect function and thus a second cohort of experimentally naïve crayfish of ~5-7cm length were exposed to 0.1, 0.5 and 1.0 M EtOH immersion.

#### Lethality:

The results of the Light/Dark transfer in the naïve ~5-7cm groups revealed substantial and consistent intoxication after 1.0 M EtOH immersion for 30 min that was not observed in the initial Open Field study. To further determine the relationship between crayfish size (age) and lethal limits of EtOH exposure we conducted a study in which two smaller size groups were exposed to different EtOH concentrations to assess lethality rates.

### 2. Supplemental Methods

**S2.1. Subjects**: Crayfish (*Procambarus clarkii*), hatched in the laboratory and divided into two groups (Cohort 3 (N=11; 7 F / 4 M), Cohort 4 (N=13; 3 F/10 M), were used for the behavioral experiments. Additional individuals were used for the lethality studies. The parents of laboratory bred individuals were obtained from a local pet store. Animals were hatched into a 10 or 20 gallon communal tank from a three clutches and then separated into groups (N=3-5) of similarly sized individuals within half to one third of a tank once reaching ~5 cm in length. Tanks were equipped with a thin layer of aquarium gravel, 2-4 huts and hideouts, free floating anacharis plants and a continuously running filter. During the course of any repeated-measures studies, nail polish markings on the carapace were used to identify individuals.

**S2.2 Behavioral assessment:** Locomotor behavior was measured in the aquatic open field and Light/Dark arenas using ANY-Maze behavior tracking of video recordings as described in the main report).

**S2.3 Ethanol Immersion exposure:** Immersion conditions were conducted as described in the main report and included ethanol (0.05, 0.1, 0.5, 1.0 M) for 30 minutes, as detailed for specific experiments below. For the behavioral studies, individuals were placed in the Center Zone of the arena after immersion and the tracking interval was initiated.

**S2.4 Supplemental Experiments**

*S2.4.1 Experiment S1: Light/Dark Transfer*

Cohort 3 (N=11; 7 F / 4 M) were ~5-7 cm in length and naïve to the arena and EtOH immersion prior to this study. All animals first received one baseline (i.e., no immersion) session in the Light/Dark arena and then were exposed to water or EtOH (0.05, 0.1 or 1.0 M) for 30 minutes, with the exposure conditions conducted in a counter balanced order. Experimental sessions were run no more frequently than twice per week (3-4 day interval). The cover placement over front vs. rear sections of the arena was balanced across individuals for this study. Out of the total sample, 2 females and 1 male completed only 1-2 conditions each due to molting and/or mortality and were omitted from the analysis.

Cohort 4 (N=13; 3 F/10 M) were ~5-7 cm in length and naïve to the arena and EtOH immersion prior to this study. All animals first received one baseline (i.e., no immersion) session in the Light/Dark arena and then were exposed to water or EtOH (0.1, 0.5 or 1.0 M) for 30 minutes, with the exposure conditions conducted in a counter balanced order. Experimental sessions were run no more frequently than twice per week (3-4 day interval). The cover placement over front vs. rear sections of the arena was balanced across individuals for this study.

*S2.4.2 Experiment S2: Lethality*

Crayfish were immersed in 0.1, 0.5 and 1.0 M EtOH for 30 minutes and then placed in normal aquarium water in a 20 cm X 20 cm plastic bin for observation. Two age groups (<2.5 cm / 5-6 weeks of age; ~2.5-4.5 cm / ~9 weeks) were assessed at each concentration with a sample size of 10 individuals per cohort. One cohort was run for most conditions save that two cohorts were run for the 2.5-4.5cm 0.5 M and the 2.5cm / 1.0 M conditions. Lethality was recorded 15 minutes into immersion, at the end of immersion, and at 30 minute intervals up to 2 h after the beginning of immersion. All of the individuals below 5 cm were euthanized at the conclusion of this experiment. Any individuals who were observed to be killed by a conspecific or that molted during the assessment interval were removed from the analysis, this involved 1 <2.5cm individual for 0.5 M and 1.0 M EtOH immersion, and two 2.5-4.5 individuals for 0.5 M EtOH immersion. Survival data for ~5-7 cm / ~11-12 weeks individuals is derived from the initial behavioral studies (i.e. those described in the main report) and reflects only those individuals exposed for 30 minutes to the respective EtOH concentration as their first-ever EtOH condition. This included 10 individuals at 1.0 M EtOH immersion, 2 individuals at 0.5 M EtOH immersion and 3 individuals at 0.1 M EtOH immersion.

**S2.5 Data analysis:**

ANY-maze was used to score time and distance in two zones of the Open Field arena, the Center versus the surrounding Periphery zone. Dependent measures included Distance Traveled, Percent of the Time spent Immobile, Speed, Center Entries, and Time spent in Center versus Periphery of the arena, divided into male and female subgroups. Treatment of missing data (e.g., due to an animal failing to enter a zone at all, failing to ever stop moving in a zone, etc) was handled by using mixed-effects analysis. The analysis strategy included a within-subjects factors for Immersion condition and the between-subjects factor of Sex.

In the Light/Dark experiments, ANY-Maze was used to score Speed and Distance in the Center, Periphery and Transition Zones as well as Entries and Time spent in all four zones. The analysis strategy included within-subjects factors for Immersion condition and Zone. To further explicate the time course of effects, the Distance and Speed measures, collapsed across zone, were analyzed by 5 minute intervals (bins) within the session.  Any significant effects for the Light/Dark experiments were followed with post-hoc analysis using Tukey correction for all multi-level, and Sidak correction for any two-level, comparisons. Prism 9 for Windows (v. 9.1.1; GraphPad Software, Inc, San Diego CA) was used for all analyses.

### S3. Supplemental Results

**S3.1 Sex Differences:** Follow-up analysis of the immersion duration study (Main report: Section 3.2) was conducted to determine potential effects of crayfish sex on Open Field behavior. There were no significant effects of Sex or the interaction of Sex with Immersion condition confirmed for Distance Traveled (**Figure S1**), Speed (**Figure S2**) or Time Immobile (**Figure S3**) in the whole arena, and for Center Entries (**Figure S4**). Similarly, there were no significant effects of Sex or the interaction of Sex with Immersion Condition confirmed for Distance Traveled, Speed or Percent Time Immobile in the Center or Periphery Zones. There was a significant effect of Sex on Center Time [$F(1, 11) = 4.94$; $P<0.05$] (**Figure S4**), however there was no significant effect of Immersion Condition or the interaction of Sex with Immersion condition confirmed for any of the measures.

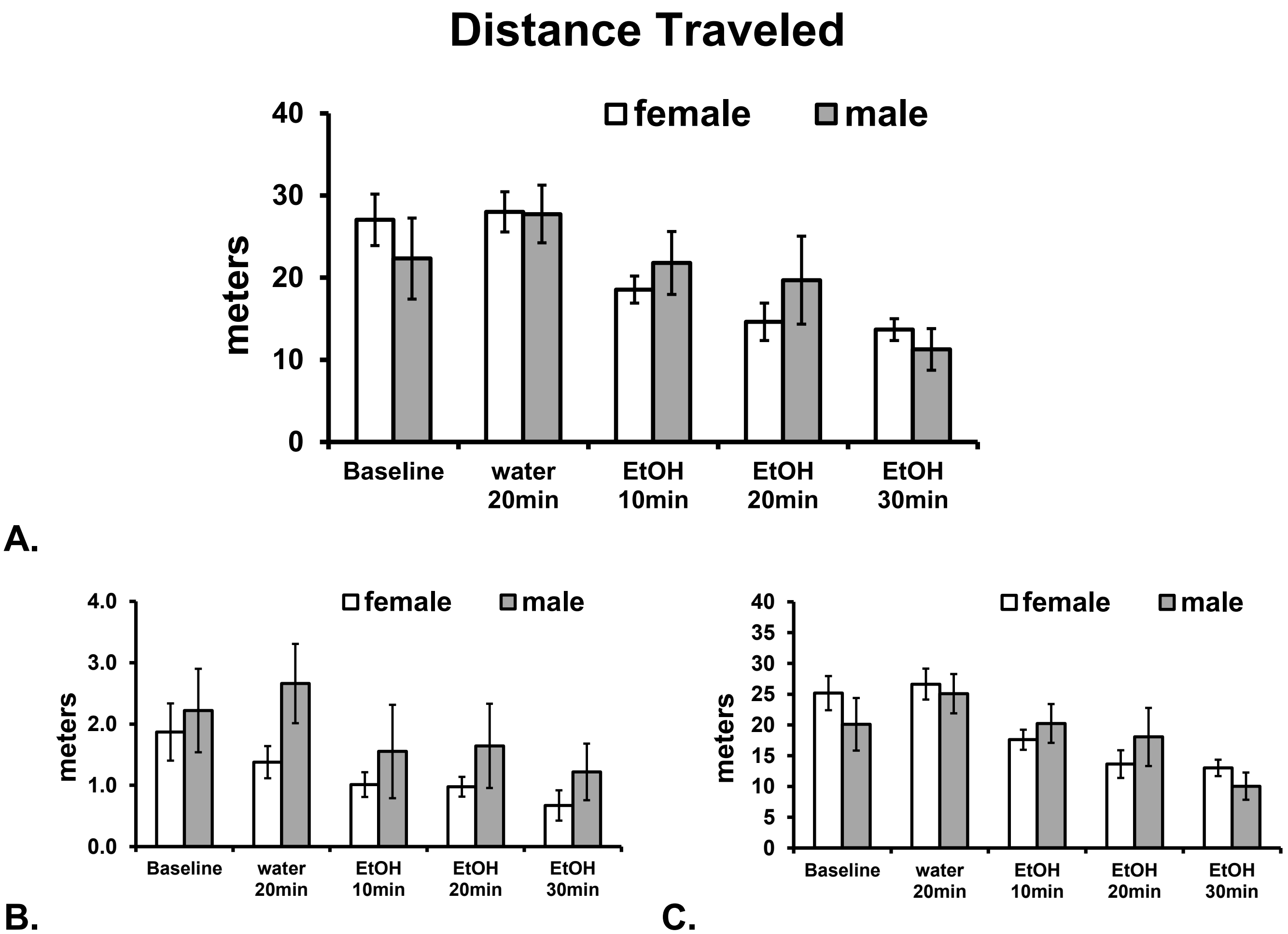


***Figure S1.*** *Mean (±SEM) Distance traveled A) overall, and in the B) Center Zone and C) Periphery Zone for female (N=7) and male (N=6) crayfish.*

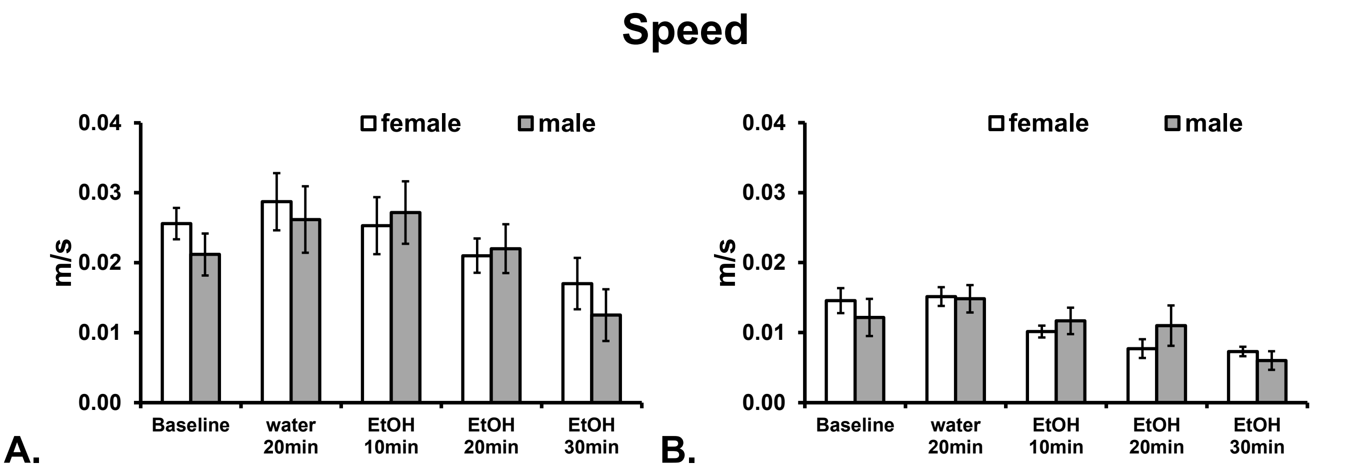


***Figure S2.*** *Mean (±SEM) Speed of movement A) in the Center Zone and B) in the Periphery Zone for female (N=7) and male (N=6) crayfish.*

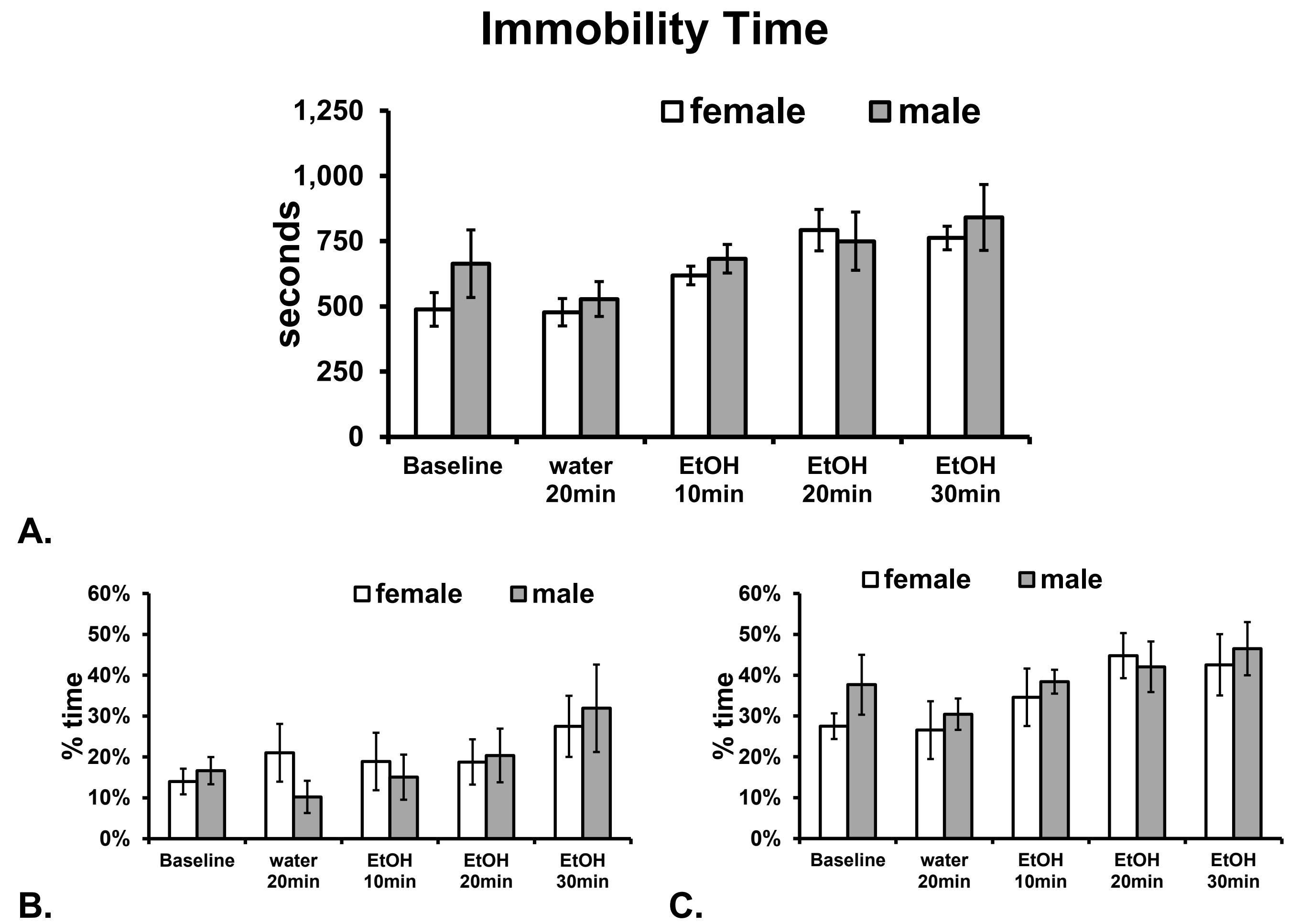


***Figure S3.*** *Mean (±SEM) A) overall time immobile, and percent time immobile in the B) Center Zone and C) Periphery Zone for female (N=7) and male (N=6) crayfish.*

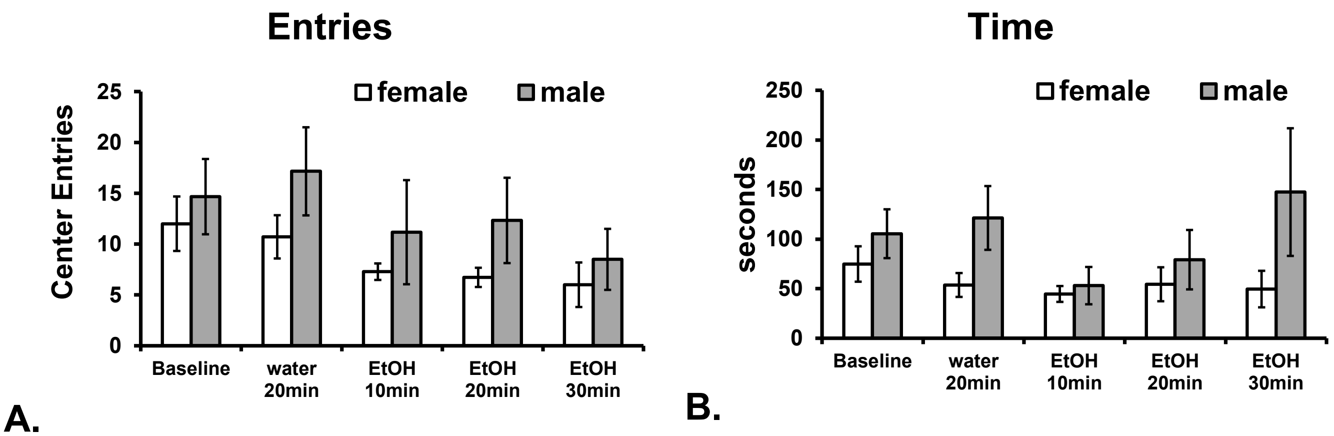


***Figure S4.*** *Mean (±SEM) A) Center Entries and B) Center Time for female (N=7) and male (N=6) crayfish.*

**S3.2 Effects of Ethanol Immersion on Light/Dark Transfer Behavior**

***<u>Cohort 3:</u>***

A group (N=8, 5 Female) of crayfish that were experimentally naïve prior to this study, were evaluated for activity in the Light/Dark arena starting with one baseline session (no prior treatment) for habituation to the arena. The group was N=10 total but two individuals only completed 1-2 conditions each and were omitted from the analysis. Due to what appeared to be a potentially stimulatory effect of 0.1 M immersion for 30 minutes in the Open Field study (Main report: Section 3.3), the immersion conditions were Water or EtOH (0.1 M, 0.5 M or 1.0 M) for 30 minutes prior to individual sessions, with conditions tested in a counter-balanced order. The behavior of this group was altered after the 1.0 M EtOH condition but not in the lower concentration conditions (**Figure S5**). Significant reductions in speed were observed in all three uncovered zones, and reduction in distance traveled in the Periphery and Transition zones. This was associated with a large increase in time spent in the Center zone and a reduction in time spent under the Cover. In essence the animals were immobilized for much of the session and did not move from where they were placed in the Center to start the test. The statistical analyses confirmed significant effects of Zone, Immersion Condition and/or the interaction for <u>Speed</u> [Zone: $F(2, 14) = 41.08$, $P<0.0001$; Immersion: $F(3, 21) = 36.34$, $P<0.0001$; Interaction: $F(6, 33) = 3.92$, $P<0.005$], <u>Time</u> [Zone: $F(3, 114) = 45.00$, $P<0.0001$; Immersion: n.s.; Interaction: $F(9, 114) = 14.93$, $P<0.0001$], <u>Distance</u> [Zone: $F(2, 14) = 48.93$, $P<0.0001$; Immersion: $F(3, 21) = 6.30$, $P<0.005$; Interaction: $F(6, 42) = 8.86$, $P<0.0001$] and <u>Entries</u> [$F(3, 24) = 11.24$, $P<0.0001$; Immersion: n.s.; Interaction: n.s.].

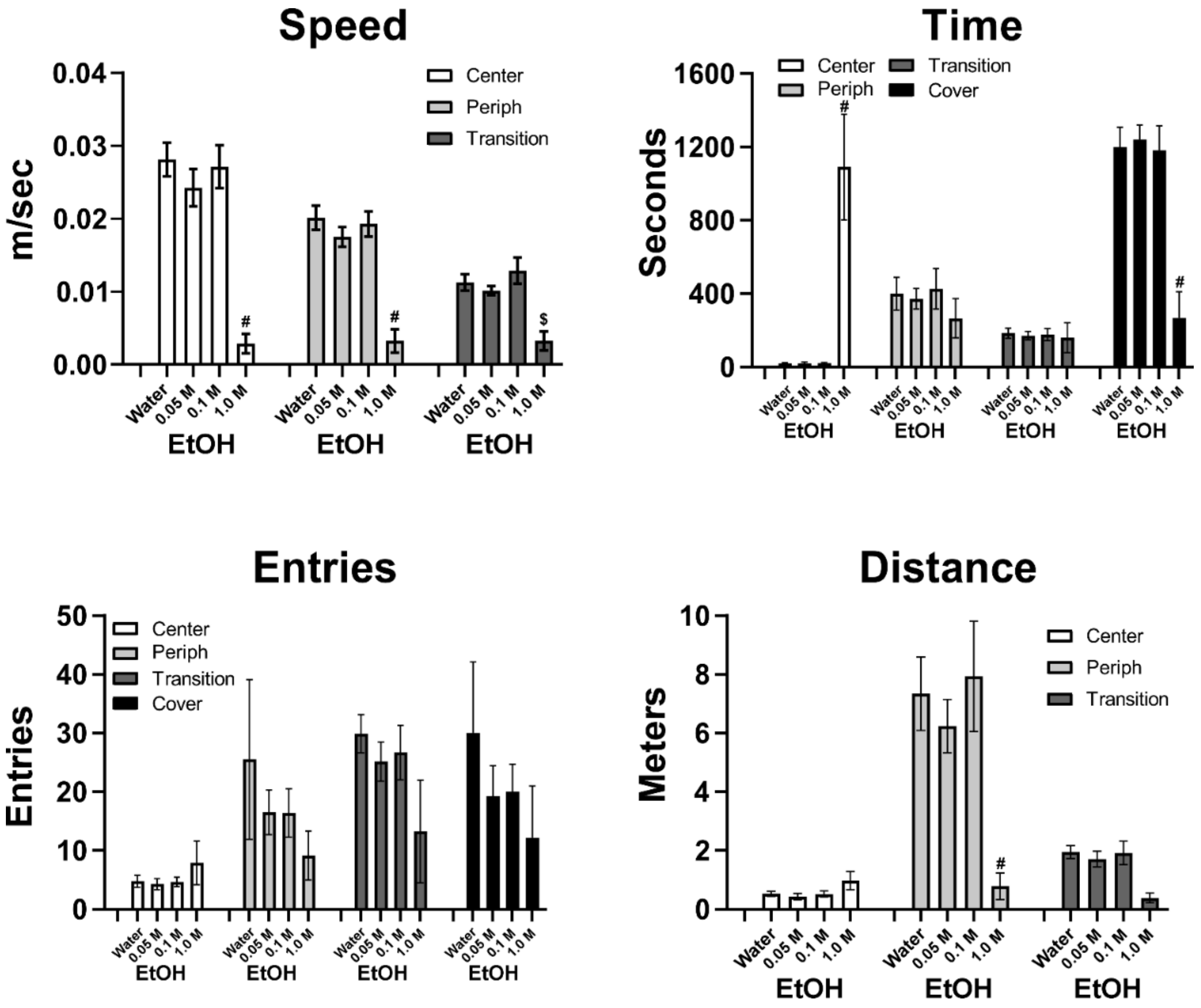


***Figure S5.*** *Mean (±SEM) Light/Dark performance for Cohort 3: The effect of 0.05-1.0 M EtOH immersion: A significant difference from all other immersion conditions, within zone, is indicated with #, and from the water and 0.1 EtOH immersion with $.*

***<u>Cohort 4:</u>***

A total of 13 individuals (3 female) from Cohort 4 were evaluated for activity in the Light/Dark arena starting with one baseline session (no prior treatment) for habituation to the arena. Crayfish were then tested for activity following 30 minutes of immersion in Water or EtOH (0.1, 0.5 or 1.0 M) prior to individual sessions in a randomized order. Individuals were lost to the study (generally due to molting) prior to completing water and 0.1 M (M97), 0.1 and 0.5 M (M92) and prior to completing 0.5M (M95). Thus, sample size for analysis cells ranged from N=11 to N=13 and a mixed-effects analysis was selected to accommodate missing cells. EtOH immersion produced dose-dependent effects on movement Speed, Time in Zone, Zone entries and Distance traveled, as shown in (**Figure S6**).

The analyses confirmed significant effects of Zone, Immersion Condition and/or the interaction for Speed [Zone: F (2, 24) = 40.88, P<0.0001; Immersion: F (3, 36) = 19.78, P<0.0001; Interaction: F (6, 38) = 5.023, P<0.001], Time [Zone: F (3, 172) = 83.92, P<0.0001; Immersion: n.s.; Interaction: F (9, 172) = 9.479, P<0.0001], Distance [Zone: F (2, 24) = 66.68, P<0.0001; Immersion: F (3, 36) = 3.570, P<0.05; Interaction: F (6, 57) = 5.083, P<0.0005] and Entries [Zone: F (3, 36) = 95.00, P<0.0001; Immersion: n.s.; Interaction: F (9, 88) = 6.846, P<0.0001].

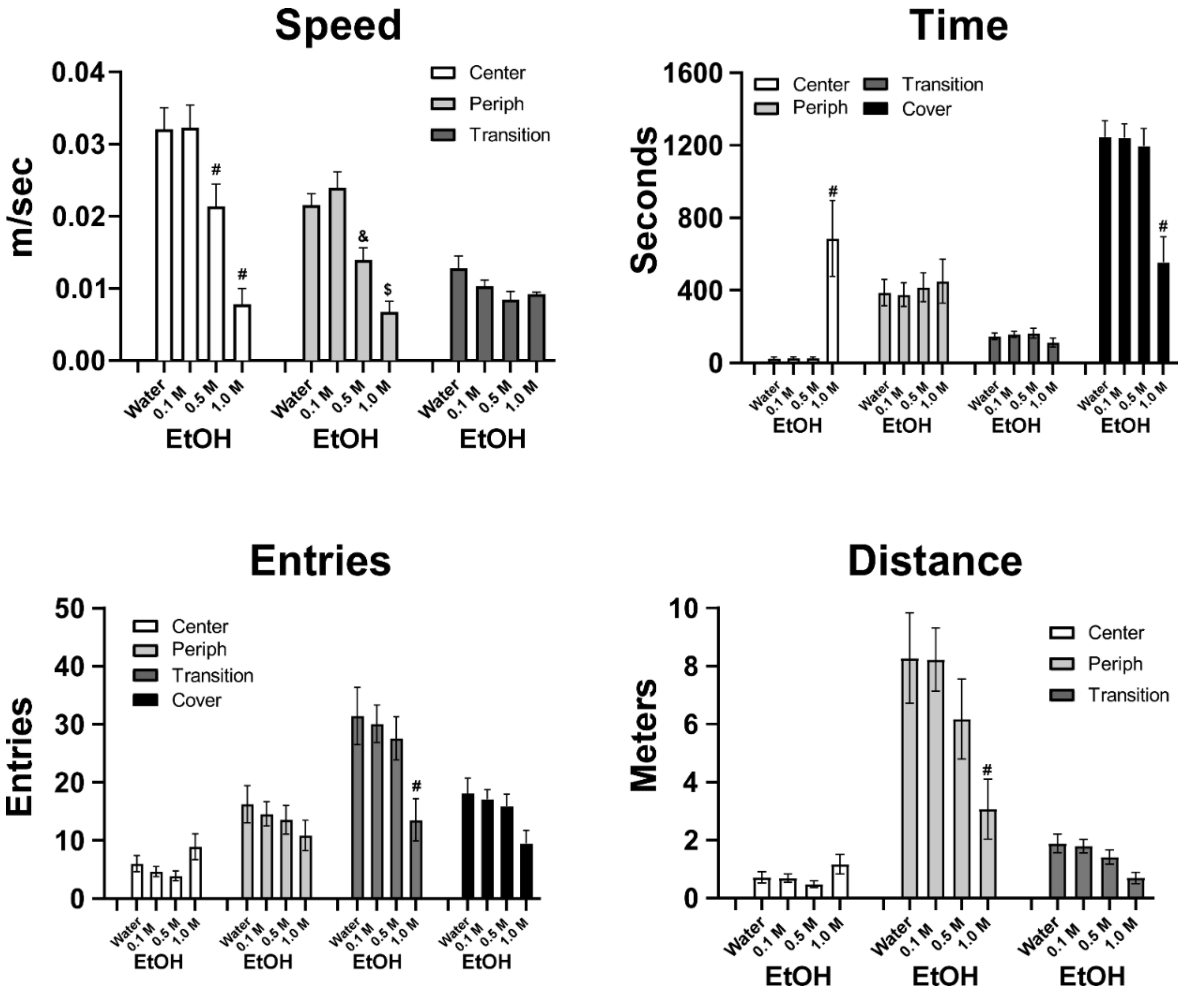


*Figure S6. The effect of 0.1-1.0 M EtOH immersion on locomotor behavior in Light/Dark Cohort 4. A significant difference from all other immersion conditions, within zone, is indicated with #, from the water and 0.1 M EtOH immersion with $ and from the 0.1 M EtOH immersion with &.*

To further explicate the time course of effects the Distance and Speed measures, collapsed across zone, patterns were analyzed by 5-minute bins within the session (**Figure S7**). The mixed-effect analyses confirmed significant effects of Time Bin, Immersion Condition and/or the interaction for

Distance [Time Bin: F (5, 60) = 13.34, P<0.0001; Immersion: F (3, 36) = 3.574, P<0.05; Interaction: F (15, 150) = 4.229, P<0.001] and for Speed [Time Bin: F (5, 60) = 15.68, P<0.0001; Immersion: F (3, 36) = 3.651, P<0.05; Interaction: F (15, 150) = 3.952, P<0.001]. The post-hoc test confirmed that crayfish traveled significantly less distance and moved significantly more slowly in the first 5 minutes after EtOH 0.5 or 1.0 M immersion compared with water or 0.1 M immersion and in the 10-minute Bin after EtOH 1.0 M immersion compared with water or 0.1 M immersion.

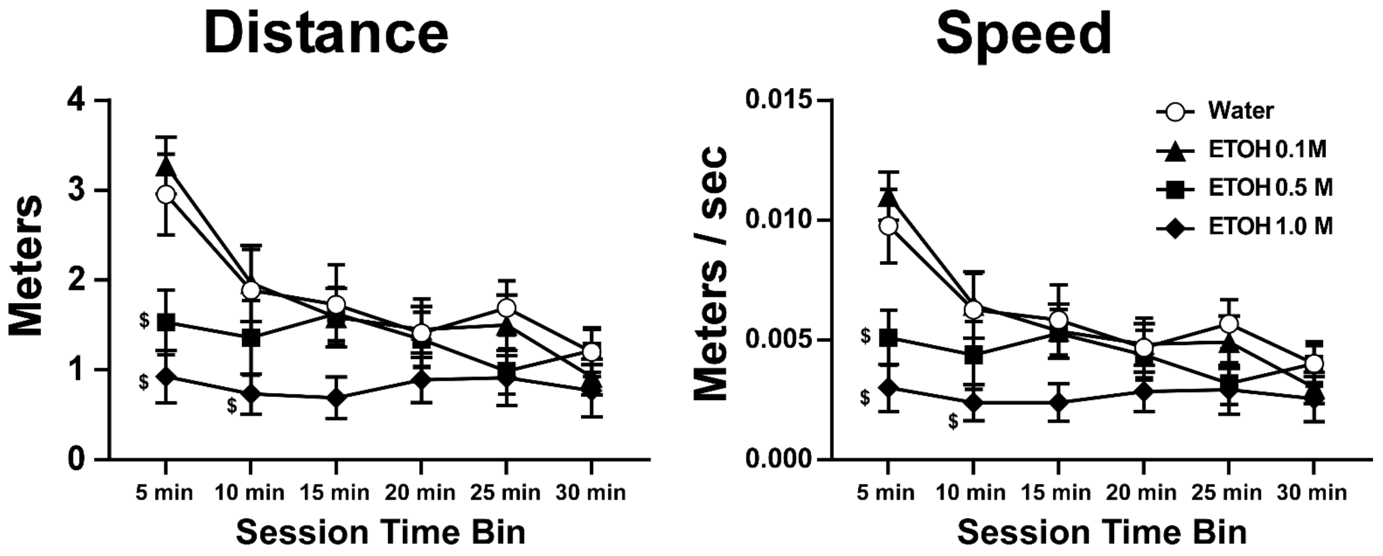


*Figure S7: The effect of 0.1-1.0 M EtOH immersion on Distance and the Speed (across all visible zones) is presented for each 5 min bin within the 30 minute session to show the time-course of effects. A significant difference from the water and 0.1 M EtOH immersion conditions is indicated with $.*

#### S3.3 Ethanol Lethality by Age

The effects of 30-minute immersion in 1.0 M EtOH were more severe in the supplemental cohorts compared with cohorts described in the main report. This was potentially because the first cohort was exposed to various durations to 1.0 M and thus only a subset of individuals experienced that dose first. Therefore, it was possible that a degree of tolerance affected the estimate of lethality. The result in the first active EtOH exposure in the repeated-immersion study, however, contradicts this explanation. And indeed, a review of the behavioral cohorts that were in the 5-7 cm range for the individuals that experience 1.0 M EtOH as their first condition found no lethality (**Figure S8**). Nevertheless, we exposed groups of juveniles of smaller size ranges to 30 minute of 0.5 M and 1.0 M EtOH to assess lethality. The 0.1 M EtOH resulted in no mortality of the <2.5 cm crayfish, however some mortality was observed after 0.5 and 1.0 M immersion. This appeared to be dose dependent, particularly in the 2.5-4.5 size range.

### S4. Supplemental Discussion

Sex Differences in the Open Field:

We show here that no significant sex differences were observed in terms of baseline behavior or in the effects of ethanol on locomotor behavior. This combines with the light/dark validation study in the main report to suggest sex differences are absent for this crayfish assay. In addition, the sex analysis can be viewed as a replication of the overall effects within subgroups.

Light/Dark Behavior in Cohorts 3 and 4:

There was no clear evidence of an anxiolytic or anxiogenic effect of EtOH immersion in the Light/Dark study with these cohorts. Although there was less time spent under the Cover after 1.0 M EtOH immersion this was attended by significantly more time spent in the Center, significantly slower Speed and reduced Distance traveled in the periphery. It was confirmed by visual inspection of videos that animals were simply immobilized in the Center after the highest dose condition. This was observed rarely in the Cohorts reported in the main report. One possibility is that although the size range in precedent literature (Swierzbinski et al., 2017) was described as “5-7 cm”, it is possible that there are significant developmental effects across this size range. We used this definition of the target size distribution, i.e., between 5 cm and 7 cm, in these cohorts. In contrast, the naïve animals used in the Cohort 1 Open field exposure duration study were mostly at the 7 cm size or slightly larger. Thus for the Light/Dark repeated dosing study we ensured a minimum of 7cm size for those investigations.

This was not designed to provide developmental information on the locomotor effects of ethanol immersion, given the sequential study in the same animals and no attempt to define developmental age beyond length and approximate age. Nevertheless, the

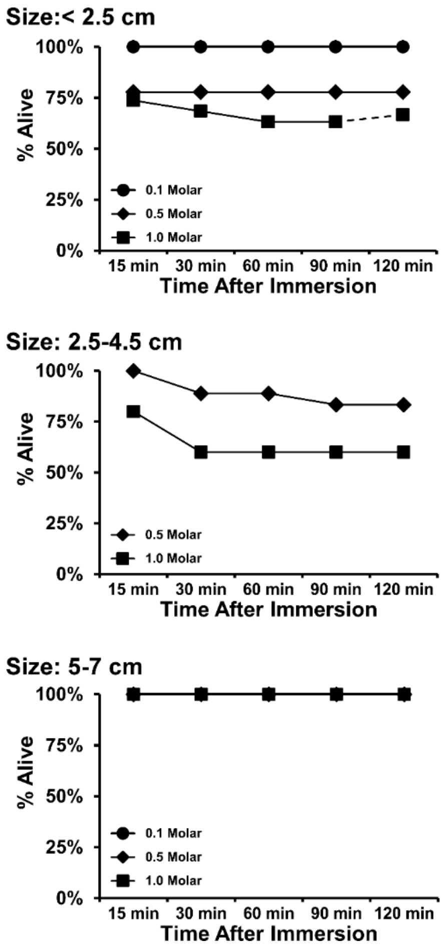


***Figure S8**: Survival after 30-minute immersion in EtOH (0.1, 0.5, 1.0 M) for A) juveniles under 2.5 cm, B) juveniles between 2.5 and 4.5 cm and C) juveniles of 5-7 cm. The 0.1 M EtOH immersion condition was omitted for the 2.5-4.5cm group. One (N=10) cohort of <2.5cm animals was only observed for 90 minutes, the second (N=9 cohort was observed for 120 minutes; thus the final point reflects the percentage survival out of 9.*

effect of 1.0 M EtOH immersion for 30 minutes was nearly identical across the first two experiments when animals had grown from late juvenile to early adult length.
